# Resilience to Endoplasmic Reticulum Stress Mitigates Calcium-Dependent Membrane Hyperexcitability Underlying Late Disease Onset in SCA6

**DOI:** 10.1101/2025.01.27.635103

**Authors:** Haoran Huang, Taylor L. Charron, Min Fu, Miranda Dunn, Deborah M. Jones, Praveen Kumar, Ashwinikumar Kulkarni, Genevieve Konopka, Vikram G. Shakkottai

## Abstract

An enduring puzzle in many inherited neurological disorders is the late onset of symptoms despite expression of function-impairing mutant protein early in life. We examined the basis for onset of impairment in Spinocerebellar ataxia type 6 (SCA6), a canonical late-onset neurodegenerative ataxia which results from a polyglutamine expansion in the voltage gated calcium channel, Cav2.1. Cerebellar Purkinje cell spiking abnormalities are seen concurrent with motor impairment in SCA6 mice but the basis for these changes in spiking is unknown. We identify endoplasmic reticulum (ER) calcium depletion as the cause for Purkinje cell spiking abnormalities and that the impairments in Purkinje cell spiking are unrelated to Cav2.1 ion-flux function. Further, intact inhibitory neurotransmission in the cerebellar cortex is necessary for Purkinje neurons to exhibit spiking abnormalities in SCA6 mice. Based on serial cerebellar transcriptome analysis, we define a mechanism of disease that is related to ER stress. Further, our studies support a model whereby proteotoxicity from misfolded mutant Cav2.1 is mitigated by a HSP90-dependent unfolded protein response (UPR) and that age-related breakdown of this response causes motor dysfunction and aberrant Purkinje cell spiking. Redundant pathways of the UPR mediate this resilience to ER stress. These studies elucidate a mechanism of resilience connecting aberrant proteostasis and calcium-dependent intrinsic membrane hyperexcitability to explain delayed disease onset more widely in age-dependent neurodegenerative disease.

**Significance Statement:** Advancing age is the single most important risk factor for neurodegenerative disease. Understanding how age intersects with genetic risk is therefore a critical challenge for neurodegenerative disease research. SCA6, a canonical late-onset degenerative cerebellar ataxia, results from a polyQ expansion in the voltage gated calcium channel, Cav2.1, encoded by *CACNA1A*. We define a mechanism of disease in SCA6 that is related to ER stress and unrelated to impaired calcium flux function of Cav2.1. Age-related decompensation of a HSP90-dependent unfolded protein response leads to disease onset. Mutant Cav2.1 misfolding as the basis for disease in SCA6 provides insight into a novel role for channelopathies to behave as proteinopathies and helps understand resilience to proteotoxicity more widely in adult-onset neurodegenerative disease.

## Introduction

Spinocerebellar ataxia type 6 (SCA6), a dominantly inherited cerebellar ataxia, is a canonical late-onset neurodegenerative disorder with symptom onset in mid- to late-life. Median age of disease onset in SCA6 is in the early-50s with some individuals who do not manifest symptoms until their 70s (1). SCA6 results from the expansion of a glutamine-encoding CAG repeat expansion in the *CACNA1A* gene, which encodes the P/Q-type voltage-gated calcium channel, Cav2.1 (2). Cav2.1 is vital for neurotransmission at the neuromuscular junction and for cerebellar Purkinje neuron function. Loss-of-function mutations in the channel cause early onset episodic ataxia or a congenital ataxia in humans (3–5). *Cacna1a* knockout mice exhibit dystonia and ataxia apparent by postnatal day 10, and die by 3-4 weeks of age (6), demonstrating the importance of the channel for cerebellar function early in life. It is therefore puzzling that symptom onset in SCA6 is late in life, in spite of the importance of Cav2.1 in early postnatal life.

SCA6^84Q/+^ mice with a hyperexpanded CAG repeat knocked into one of the endogenous mouse *Cacna1a* alleles have been generated to model human SCA6 (7). Heterozygous SCA6^84Q/+^ mice, the most precise genetic model currently available, develop motor impairment at 19 months of age, in a manner similar to late symptom-onset in individuals with SCA6. Motor impairment in SCA6^84Q/+^ mice is concurrent with abnormally irregular spiking of cerebellar Purkinje neurons. The irregular spiking is the likely driver of motor impairment, as improving spiking regularity pharmacologically, also improves motor dysfunction (8). The relationship between the CAG repeat expansion in *Cacna1a* and alterations in spiking is unclear.

Purkinje neurons, the sole output from the cerebellar cortex, are autonomous pacemakers that fire action potentials at a high frequency even in the absence of synaptic input (9, 10). Purkinje neuron pacemaking is possible through the interplay of a specific group of ion channels, whose biophysical properties allow for fine tuning of the frequency and regularity of spiking (10). In a number of mouse models of spinocerebellar ataxia (SCA) that result from a CAG repeat expansion in the respective genes, a reduction in firing frequency or spiking regularity are seen concurrent with onset of motor impairment (8, 11–17). A shared reduction in ion channel transcript levels of *Kcnma1*, *Cacna1g*, *Trpc3* and *Itpr1* is seen in SCA1, SCA2, and SCA7 (13, 15, 16, 18). Although SCA6^84Q/+^ mice display alterations in spiking similar to that seen in models of SCA1 and SCA7, the basis for the changes in spiking remains unexplored.

The endoplasmic reticulum (ER) is vital for proper protein folding. Protein folding in the ER is mediated through a number of resident chaperone proteins. The ER is also the major site of calcium storage in the cell. Perturbation of ER-luminal calcium concentrations inhibits chaperone function (19). A number of conditions, including protein misfolding and ER calcium depletion, result in ER stress. Cells activate a signaling pathway known as the unfolded protein response (UPR) to alleviate ER stress. There are three canonical pathways that have been identified, including XBP1, ATF6, and PERK-eIF2α (20). These pathways are thought to separately mediate ER stress responses under conditions of abnormal proteostasis. *Xbp1* transcripts undergo unconventional splicing by IRE1α, which leads to generation of a transcription factor sXBP1 (21). Increased function of XBP1 has been linked to enhanced chaperone-mediated protein folding among other functions related to the UPR (22, 23). Upon ER stress, ATF6 translocates to the Golgi apparatus where it is cleaved to generate ATF6f. ATF6f subsequently induces expression of *Xbp1* and regulates the UPR together with XBP1 (24, 25). PERK activation induces phosphorylation of eIF2α (peIF2α), which inhibits protein synthesis to prevent ER protein overload (20). Under conditions of unmitigated ER stress, eIF2α phosphorylation tips cells towards apoptosis (26). Glutamine encoding CAG repeat (polyQ) disorders are postulated to result in misfolded proteins, and activate ER stress responses (27). However, whether and how ER stress leads to disease in polyQ ataxia remains unknown.

We examined the basis for the late-onset disease phenotype in SCA6^84Q/+^ mice, where motor impairment is only evident at 19 months. Utilizing unbiased transcriptome analysis in presymptomatic SCA6^84Q/+^ mice, we discovered a homeostatic response that engages the UPR at 6 months, long preceding onset of motor impairment. This resilience to ER stress is associated with activation of all three canonical UPR pathways and mediated through the HSP90 chaperone machinery. ER-stress mediated activation of a calcium-release activated calcium current drives the increased membrane excitability and aberrant spiking in SCA6^84Q/+^ Purkinje neurons. We have identified a mechanism for preservation of Purkinje neuron health in SCA6 that explains the late onset of the disease phenotype, and the mechanism for cerebellar impairment in SCA6.

## Results

### Irregular Purkinje neuron spiking in SCA6 is not due to ion channel transcript dysregulation

SCA6^84Q/+^ mice, where a hyperexpanded CAG encoding polyglutamine (polyQ) repeat is knocked into one of the endogenous mouse *Cacna1a* alleles, represent a genetically precise model of the dominantly inherited human neurological disorder. Heterozygous SCA6^84Q/+^ mice do not display motor impairment until 19 months of age (7, 8). In other models of SCA, motor dysfunction is associated with irregular cerebellar Purkinje neuron spiking (13, 28). Prior work has identified Purkinje neuron spiking irregularity in 19-month SCA6^84Q/+^ mice (8). It is unclear, however, whether asynchronous onset of irregular Purkinje neurons spiking reaches a threshold at 19 months to cause motor dysfunction. To examine Purkinje neuron spiking regularity prior to the onset of motor impairment, we performed noninvasive cell-attached patch-clamp recordings in acute cerebellar slices in SCA6^84Q/+^ mice at near physiological temperatures at various ages. Purkinje neurons showed no change in spiking regularity or firing frequency in SCA6^84Q/+^ mice at 3 months (Figure 1A-C, S1A), 6 months (Figure 1D-F, S1B), or 12 months (Figure 1G-I, S1C) of age. Significant spiking irregularly was observed only in 19-month SCA6^84Q/+^ mice compared to wild-type (WT) mice (Figure 1J-L), with no alteration in firing frequency (Figure S1D). Both male and female 19-month SCA6^84Q/+^ mice exhibited similar Purkinje neuron spiking irregularity (data not shown), and therefore, subsequent studies were done without regard to sex. These data suggest that in SCA6^84Q/+^ mice, spiking irregularity is likely the driver of motor impairment. Alterations in ion channel transcript levels are associated with spiking impairment in SCA1 (18, 28, 29) and SCA7 (13) mouse models. To examine whether changes in the expression of ion channels may account for irregular spiking, we used quantitative RT-PCR and compared several key ion channel transcript levels in the cerebella of WT and SCA6^84Q/+^ mice at different ages (Figure S1E-H). The key ion channels transcripts examined included calcium channels (including *Cacna1g, Cacna1a, Trpc3*, and *Itpr1*), and calcium-activated potassium channels (including *Kcnma1* and *Kcnn2*). No significant changes in transcript levels of these ion channels were identified at 3 months of age (Figure S1E). Although small but significant cerebellar changes were noted in transcript levels of several ion channels in 6-month (Figure S1F), 12-month (Figure S1G), and 19-month SCA6^84Q/+^ mice (Figure S1H), the changes were inconsistent across the different ages. We further compared protein levels of Cav2.1 (encoded by *Cacna1a*) and Cav3.1 (encoded by *Cacna1g*) in WT and SCA6^84Q/+^ mice at 6 months and 19 months of age using immunostaining. The intensity of Cav2.1 and Cav3.1 staining in the cerebellar molecular layer were comparable between WT and SCA6^84Q/+^ mice at both ages (Figure S1I-J), indicating the expression of Cav2.1 and Cav3.1 are unchanged despite small changes in ion channel transcript levels. Taken together, these data suggest that age-dependent Purkinje neuron dysfunction is not caused by changes in expression of ion channels.

**Figure 1.**
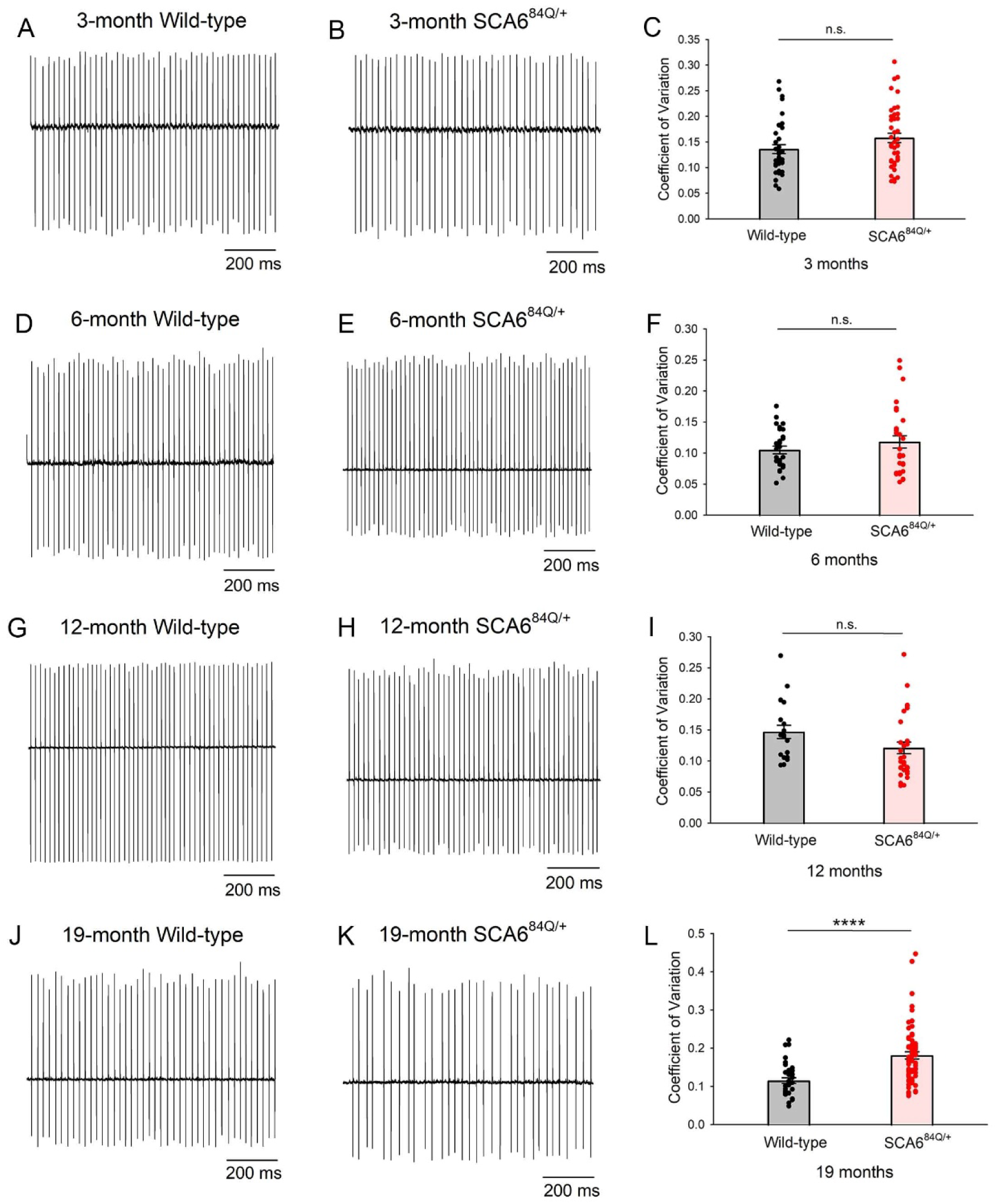
Age-dependent irregular Purkinje neuron spiking in SCA6 is not due to changes in action potential shape. A) Representative trace of spontaneous Purkinje neuron spiking from acute cerebellar slices of 3-month WT mice. B) Representative trace of spontaneous Purkinje neuron spiking from acute cerebellar slices of 3-month SCA6^84Q/+^ mice. C) Purkinje neuron spiking regularity, indicated by the coefficient of variation of the interspike interval, is comparable between 3-month WT (N = 3) and SCA6^84Q/+^ mice (N = 3). WT cells: n = 37, SCA6^84Q/+^ cells: n = 42. Student’s t-test, n.s.: Not significant. D) Representative trace of spontaneous Purkinje neuron spiking from acute cerebellar slices of 6-month WT mice. E) Representative trace of spontaneous Purkinje neuron spiking from acute cerebellar slices of 6-month SCA6^84Q/+^ mice. F) Purkinje neuron spiking regularity, indicated by the coefficient of variation of the interspike interval, is comparable between 6-month WT (N = 3) and SCA6^84Q/+^ mice (N = 3). WT cells: n = 28, SCA6^84Q/+^ cells: n = 30. Student’s t-test, n.s.: Not significant. G) Representative trace of spontaneous Purkinje neuron spiking from acute cerebellar slices of 12-month WT mice. H) Representative trace of spontaneous Purkinje neuron spiking from acute cerebellar slices of 12-month SCA6^84Q/+^ mice. I) Purkinje neuron spiking regularity, indicated by the coefficient of variation of the interspike interval, is comparable between 12-month WT (N = 4) and SCA6^84Q/+^ mice (N = 3). WT cells: n = 20, SCA6^84Q/+^ cells: n = 31. Student’s t-test, n.s.: Not significant. J) Representative trace of spontaneous Purkinje neuron spiking from acute cerebellar slices of 19-month WT mice. K) Representative trace of spontaneous Purkinje neuron spiking from acute cerebellar slices of 19-month SCA6^84Q/+^ mice showing spiking irregularity. L) The coefficient of variation of the interspike interval, representing spiking regularity, is significantly abnormal in 19-month SCA6^84Q/+^ mice (N = 11) compared to WT mice (N = 4). WT cells: n = 35, SCA6^84Q/+^ cells: n = 64. Student’s t-test, ****P < 0.0001.

### Irregular Purkinje neuron spiking in SCA6 results from altered intrinsic membrane excitability that is facilitated by intact inhibitory neurotransmission

The pacemaker firing of Purkinje neurons is modulated by synaptic input. In order to examine whether the irregular spiking observed in 19-month SCA6^84Q/+^ mice is driven by a synaptic or intrinsic mechanism, we evaluated Purkinje neuron spiking in the presence of inhibitors of fast synaptic transmission. In cerebellar slices from 19-month SCA6^84Q/+^ mice, perfusion of a combination of picrotoxin, a GABA_A_ receptor antagonist, and 6,7-dinitroquinoxaline-2,3-dione (DNQX), a non-NMDA receptor antagonist, significantly improved Purkinje neuron spiking irregularity (Figure S2A) without affecting firing frequency (Figure S2B). Unexpectedly, picrotoxin alone improved Purkinje neuron spiking irregularity (Figure 2A), whereas DNQX alone had no effect on modulating Purkinje neuron spiking irregularity (Figure S2C) in 19-month SCA6^84Q/+^ mice. Neither picrotoxin alone nor DNQX alone altered Purkinje neuron firing frequency (Figure 2B and S2D). To further determine whether alterations in inhibitory neurotransmission in Purkinje neurons in SCA6^84Q/+^ mice is responsible for irregular Purkinje neuron spiking, we examined spontaneous inhibitory post-synaptic currents (IPSCs). We measured the picrotoxin-sensitive (Figure S2E) inhibitory postsynaptic current (IPSC) frequency and regularity in Purkinje neurons of 19-month WT and SCA6^84Q/+^ mice. There was no difference in IPSC frequency (Figure 2C) or regularity (Figure 2D). These data suggest that although intact inhibitory synaptic transmission is necessary for SCA6 Purkinje neuron pathophysiology, alterations in inhibitory neurotransmission are not directly the cause of Purkinje neuron spiking irregularity.

**Figure 2.**
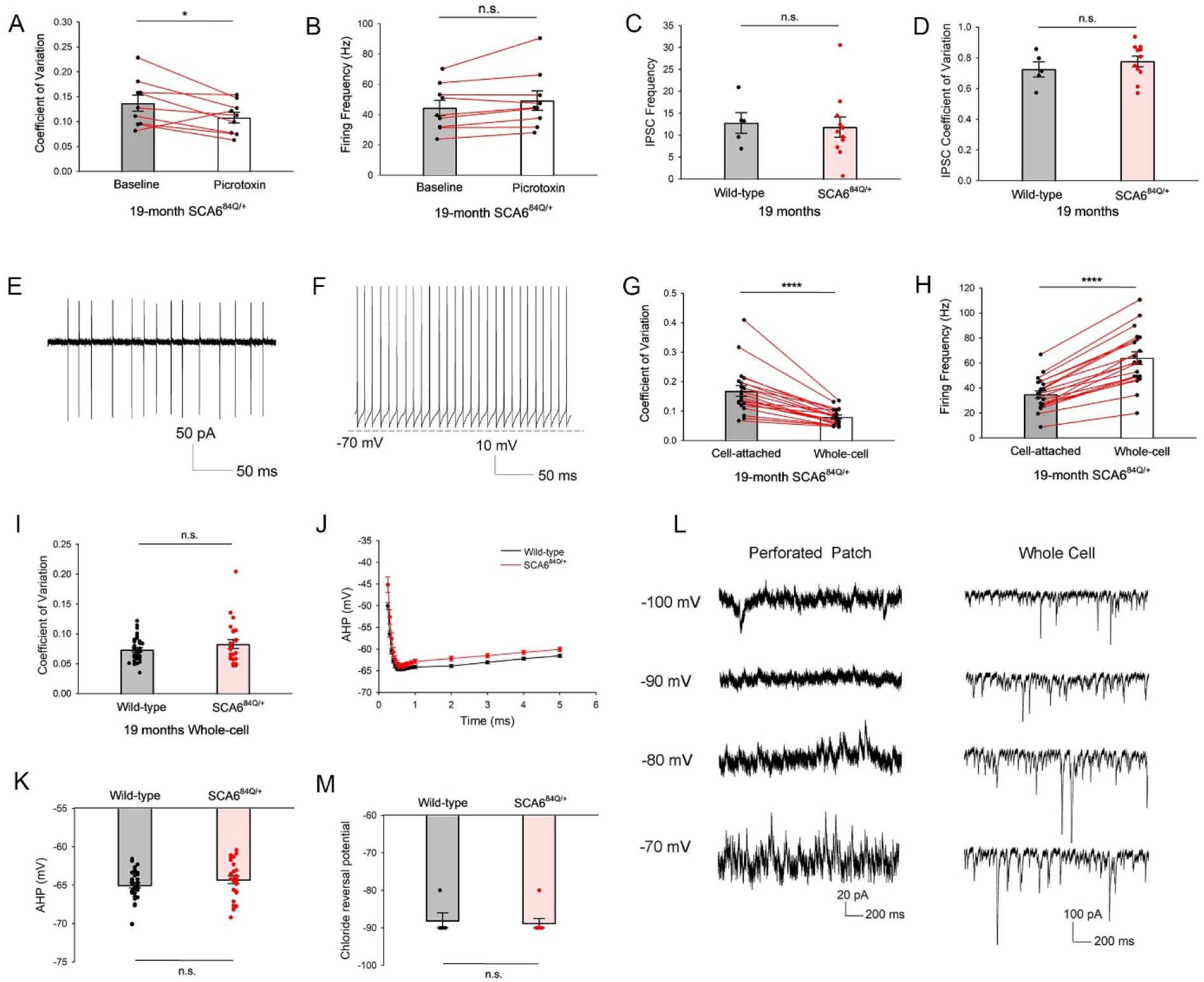
Irregular Purkinje neuron spiking in SCA6^84Q/+^ mice is due to changes in intrinsic membrane excitability. A) Picrotoxin improves Purkinje neuron spiking irregularity in 19-month SCA6^84Q/+^ mice. n = 9 cells. Paired t-test, *P < 0.05. B) Picrotoxin has no effect on Purkinje neuron firing frequency in 19-month SCA6^84Q/+^ mice. n = 9 cells. Paired t-test, n.s.: Not significant. C) Frequency of IPSCs in Purkinje neuron is comparable between 19-month WT and SCA6^84Q/+^ mice. WT cells: n = 5, SCA6^84Q/+^ cells: n = 11. Student’s t-test, n.s.: Not significant. D) Regularity of IPSCs in Purkinje neuron is comparable between 19-month WT and SCA6^84Q/+^ mice. WT cells: n = 5, SCA6^84Q/+^ cells: n = 11. Student’s t-test, n.s.: Not significant. E) Representative trace of Purkinje neuron spiking in the cell-attached configuration from 19-month SCA6^84Q/+^ mice. F) Representative trace of Purkinje neuron spiking in the whole-cell configuration from the same cell in (E). G) Patch break-in in Purkinje cells whose spiking was monitored noninvasively in the cell-attached configuration improves Purkinje neuron spiking irregularity in 19-month SCA6^84Q/+^ mice. n = 20 cells. Paired t-test, ****P < 0.0001. H) Patch break-in in Purkinje cells whose spiking was monitored noninvasively in the cell-attached configuration increases Purkinje neuron firing frequency in 19-month SCA6^84Q/+^ mice. n = 20 cells. Paired t-test, ****P < 0.0001. I) Purkinje neuron spiking regularity in the whole-cell patch clamp configuration is comparable between 19-month WT and SCA6^84Q/+^ mice. WT cells: n = 33, SCA6^84Q/+^ cells: n = 24. Student’s t-test, n.s.: Not significant. J) Characterization of the AHP in 19-month WT (n = 33) and SCA6^84Q/+^ (n = 26) Purkinje neurons. K) The amplitude of the AHP as defined as the most hyperpolarized membrane potential, is similar between WT and SCA6^84Q/+^ Purkinje neuron at 19-months. Student’s t-test, n.s.: Not significant. L) The chloride reversal potential is comparable between 19-month WT (n = 4) and SCA6^84Q/+^ Purkinje neurons (n = 4). Student’s t-test, n.s.: Not significant. M) Left: Representative traces of IPSCs using gramicidin perforated-patch recordings showing a chloride reversal potential of −90 mV in 19-month SCA6^84Q/+^ Purkinje neurons. Please note that current is inward at −100 mV and outward at −80 mV. R_s_ = 28 MΩ. Right: Representative traces showing IPSCs in 19-month SCA6^84Q/+^ Purkinje neuron in the whole-cell patch clamp configuration. Please note that the IPSCs are inward at all the voltages shown. R_s_ = 10 MΩ.

To examine whether altered intrinsic excitability may in fact be responsible for Purkinje neuron irregular spiking, we recorded spiking from the same 19-month SCA6^84Q/+^ Purkinje neurons in the cell-attached configuration and subsequently the whole-cell patch clamp configuration. If irregular Purkinje neuron spiking in SCA6^84Q/+^ mice is due to a synaptic mechanism, the spiking irregularity seen in the cell-attached configuration would be preserved in the whole-cell patch-clamp configuration in 19-month SCA6^84Q/+^ mice. We performed noninvasive cell-attached recordings and following patch break-in, whole-cell voltage recordings in the same cells (Figure 2E,F). Surprisingly, spiking irregularity was significantly improved in the whole-cell configuration compared to the cell-attached configuration in 19-month SCA6^84Q/+^ Purkinje neurons (Figure 2G). Purkinje neuron firing frequency also increased significantly in the whole-cell configuration (Figure 2H). Further, spiking regularity and firing frequency between 19-month WT and SCA6^84Q/+^ mice in the whole-cell configuration were comparable (Figure 2I,S2F), unlike in the cell-attached configuration where spiking was irregular in SCA6^84Q/+^ mice compared to wild-type controls (Figure 1L). These results suggest that an abnormality in intrinsic Purkinje neuron membrane excitability explains the irregular spiking in 19-month SCA6^84Q/+^ Purkinje neurons, and that dialysis/buffering of intracellular contents, that are in turn influenced by intact GABA_A_ neurotransmission, ameliorates the cause of the irregular spiking in SCA6^84Q/+^ Purkinje neurons.

An intrinsic mechanism that results in irregular Purkinje neuron spiking is abnormalities in the afterhyperpolarization (AHP) phase of the action potential (13, 28, 30), which is sensitive to intracellular calcium buffering. Also, the AHP is strongly influenced by calcium flux through Cav2.1 and a reduction in Cav2.1 currents produces irregular Purkinje neuron spiking (30). To determine whether irregular Purkinje neuron spiking in 19-month SCA6^84Q/+^ mice results from changes in the AHP, we compared the amplitude of the AHP during spontaneous spiking, between 19-month WT and SCA6^84Q/+^ mice using whole-cell patch-clamp recordings with intracellular calcium buffering that preserves normal Purkinje neuron AHP amplitude (13, 16, 28). No significant changes in AHP amplitude were observed (Figure 2J,K). Other spike parameters analyzed, including spike peak, depolarization slope, repolarization slope, spike half-width, and input resistance, also failed to reveal biologically meaningful differences between WT and SCA6^84Q/+^ Purkinje neurons (Table S1) that would explain the spiking irregularity. Taken together, these results suggest that the irregular Purkinje neuron spiking observed in 19-month SCA6^84Q/+^ mice result from an intrinsic mechanism different from alterations in expression or function of ion channels, including Cav2.1, previously implicated in producing this aberrant spiking phenotype.

It is puzzling that preserved GABA_A_-mediated inhibitory neurotransmission is necessary to increase Purkinje neuron intrinsic membrane excitability in SCA6^84Q/+^ Purkinje neurons. During development, GABA_A_-neurotransmission causes membrane depolarization. This in turn is mediated by elevated cytosolic chloride concentrations and the resulting more depolarized chloride reversal potential (31, 32). We measured the chloride reversal potential in 19-month WT and SCA6^84Q/+^ Purkinje neurons using gramicidin perforated-patch recordings. Gramicidin channels are impermeable to divalent cations and all anions (33), allowing us to measure the chloride reversal potential of spontaneous Purkinje neuron IPSCs. Perforated-patch recordings revealed a chloride reversal potential of −90 mV in 19-month SCA6^84Q/+^ Purkinje neurons, which is comparable to the chloride reversal potential in WT Purkinje neurons (Figure 2L,M). Taken together, we conclude that intact GABA_A_ inhibitory transmission affects intracellular contents in a way that facilitates an increase in Purkinje neuron intrinsic membrane excitability and leading to irregular spiking in SCA6 Purkinje neurons.

### Identification of ER stress as a potential driver of disease

Prior studies have demonstrated the utility of unbiased transcriptome analysis to identify the molecular basis for changes in Purkinje neuron intrinsic membrane excitability in cerebellar ataxia (13, 18). We performed bulk RNA sequencing (RNAseq) from cerebella of WT and SCA6^84Q/+^ mice at 3, 6 12 and 19 months of age and used unbiased Weighted Gene Correlation Network Analysis (WGCNA) to identify coexpressed gene networks. We then examined which gene networks were significantly enriched in differentially expressed genes (DEGs) between WT and SCA6^84Q/+^ mice. We determined whether DEGs between WT and SCA6^84Q/+^ mice at 3, 6 12 and 19 months of age could provide insight into mechanisms underlying resilience to spiking abnormalities at presymptomatic ages, and thereby also provide insight into mechanism for spiking irregularities in symptomatic 19-month SCA6^84Q/+^ mice. WGCNA identified correlated gene expression modules as a function of age (Figure 3A, top, Table S2). None of these gene expression modules displayed correlated changes as a function of genotype (SCA6^84Q/+^ versus WT). Gene expression modules containing ion channel genes implicated in inherited ataxia are contained in the white and yellow modules. The yellow module had no significant enrichment of DEGs between WT and SCA6^84Q/+^ mice at any age (Figure 3A, middle, Table S2). No ion channel gene transcripts in the white module were differentially expressed between WT and SCA6^84Q/+^ mice (Figure 3A, middle, Table S2). The gene expression module that showed the most significant enrichment of genes differentially expressed between WT and SCA6^84Q/+^ mice was the lightcyan module (highlighted in Figure 3A, bottom). The lightcyan module showed a highly significant enrichment of DEGs that were upregulated in SCA6^84Q/+^ compared to WT mice at 6 months of age. Examination of the genes within the lightcyan module revealed ER-resident UPR-related genes, with transcripts for upregulated ER chaperones (*Hsp90b1*, *Hyou1*, *Hspa5*, *Dnajb9*, *Dnajb11*, *Pdia3*, and *Dnajc3*) and ER calcium buffers (*Calr*) (Figure 3B) at 6 months of age. Among the transcripts that were significantly reduced at 12 months of age in the lightcyan module, *Creld2* is also a UPR-related gene (Figure 3B). The UPR is a protective pathway that is activated by ER stress (20). UPR-related genes are regulated by 3 major pathways, namely, XBP1, ATF6, and PERK-eIF2α. The DEGs in the lightcyan module are known transcriptional targets of XBP1 and ATF6 (34). This suggests that enhancement of UPR in response to ER stress may play a homeostatic role in SCA6^84Q/+^ mice at presymptomatic ages.

**Figure 3.**
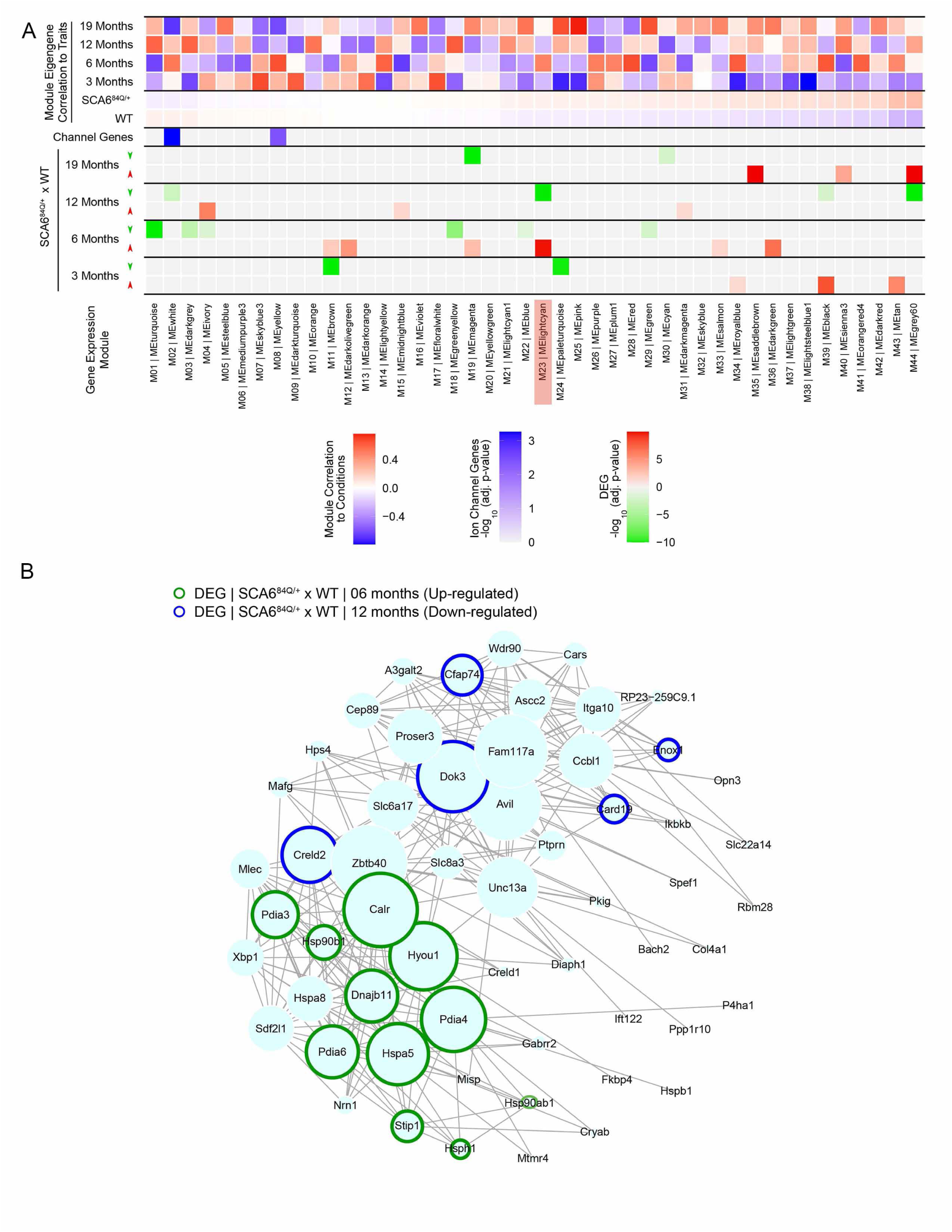
Weighted Gene Correlation Network Analysis (WGCNA) identifies ER stress as a potential driver of disease in SCA6. A) Top: WGCNA identifies correlated gene modules as a function of age but not as a function of genotype (SCA6^84Q/+^ vs WT); Middle: Ion channel genes implicated in inherited ataxia are enriched in white and yellow modules. The yellow module exhibits no enrichment of DEGs between WT and SCA6^84Q/+^ mice at any age; The ion channels gene transcripts in the white module are not differentially expressed in SCA6^84Q/+^ mice; Bottom: WGCNA identifies genes within the lightcyan module (highlighted) as significantly upregulated in 6-month SCA6^84Q/+^ compared to WT mice. B) Network plot showing the genes within the lightcyan module and the corresponding changes in expression at different ages between WT and SCA6^84Q/+^ mice. There is upregulation of ER resident chaperones (*Hsp90b1, Hyou1, Hspa5, Dnajb11*), folding enzymes (*Pdia3, Pdia4, Pdia6)* and ER calcium buffers (*Calr*) in 6-month SCA6^84Q/+^ mice. Downregulation of the UPR-related gene, *Creld2*, is noted in 12-month SCA6^84Q/+^ mice. The size of the circle represents the weight of the correlation with other module genes. Note that genes that are not outlined exhibit correlated expression with other module genes but the expression is not differentially altered in SCA6^84Q/+^ mice (upregulated DEGs outlined in green; downregulated DEGs outlined in blue).

### Alterations in all three canonical UPR pathways are present in SCA6

Since we identified UPR-related genes as the most significantly enriched DEGs between WT and SCA6^84Q/+^ mice, we sought to examine whether there are changes in proteostasis in SCA6^84Q/+^ mice. Three canonical UPR pathways are identified: XBP1, ATF6, and PERK-eIF2α (20). Of these, XBP1 is the most evolutionarily conserved UPR pathway (20). We observed that *Xbp1* is co-expressed with other UPR-related genes in the lightcyan module (Figure 3B). *Xbp1* undergoes non-canonical splicing to generate the active transcription factor sXBP1 (21). To investigate the role of XBP1 in proteostasis in pre-symptomatic SCA6^84Q/+^ mice, we examined spliced *Xbp1* (*sXbp1*) transcript levels in the cerebella of 6-month SCA6^84Q/+^ mice. We examined activation of UPR pathways separately in male and female mice. A significant elevation of *sXbp1* transcript levels was seen in both male and female SCA6^84Q/+^ mice at 6 months of age, although the degree of *sXbp1* elevation was higher in female mice (Figure 4A). We also examined the protein levels of sXBP1 in the whole cerebellum of 6-month male and female SCA6^84Q/+^ mice (Figure 4B). Paradoxically, sXBP1 protein levels were reduced in both male and female SCA6^84Q/+^ mice, although the reduction was only significant in male mice (Figure 4C). We further examined protein expression of sXBP1 using immunostaining to explore the subcellular localization of sXBP1 in Purkinje neurons (Figure 4D). sXBP1 predominantly localized to nuclei of Purkinje neurons. The intensity of sXBP1 was significantly reduced in the nuclei of 6-month SCA6^84Q/+^ Purkinje neurons (Figure S4A). Interestingly, despite decreased intensity of sXBP1 staining in most SCA6^84Q/+^ Purkinje neuron nuclei (Figure 4D), we observed more focal nuclear localization of sXBP1 in some 6-month SCA6^84Q/+^ Purkinje neurons (Figure 4D, inset). Together with our previous data showing upregulation of *sXbp1* targets in cerebella of 6-month SCA6^84Q/+^ mice, these data suggest enhanced function of sXBP1 in 6-month SCA6^84Q/+^ Purkinje neurons despite reduced total protein levels of sXBP1 in the whole cerebellum. The UPR is canonically mediated by two other pathways in addition to XBP1; namely ATF6 and PERK-eIF2α (20, 35). Cerebellar ATF6 levels were significantly reduced in 6-month male but not female SCA6^84Q/+^ mice (Figure 4E,F). Cerebellar peIF2α relative to total eIF2α (peIF2α/eIF2α) levels were unchanged in both 6-month male and female SCA6^84Q/+^ mice (Figure 4G,H).

**Figure 4.**
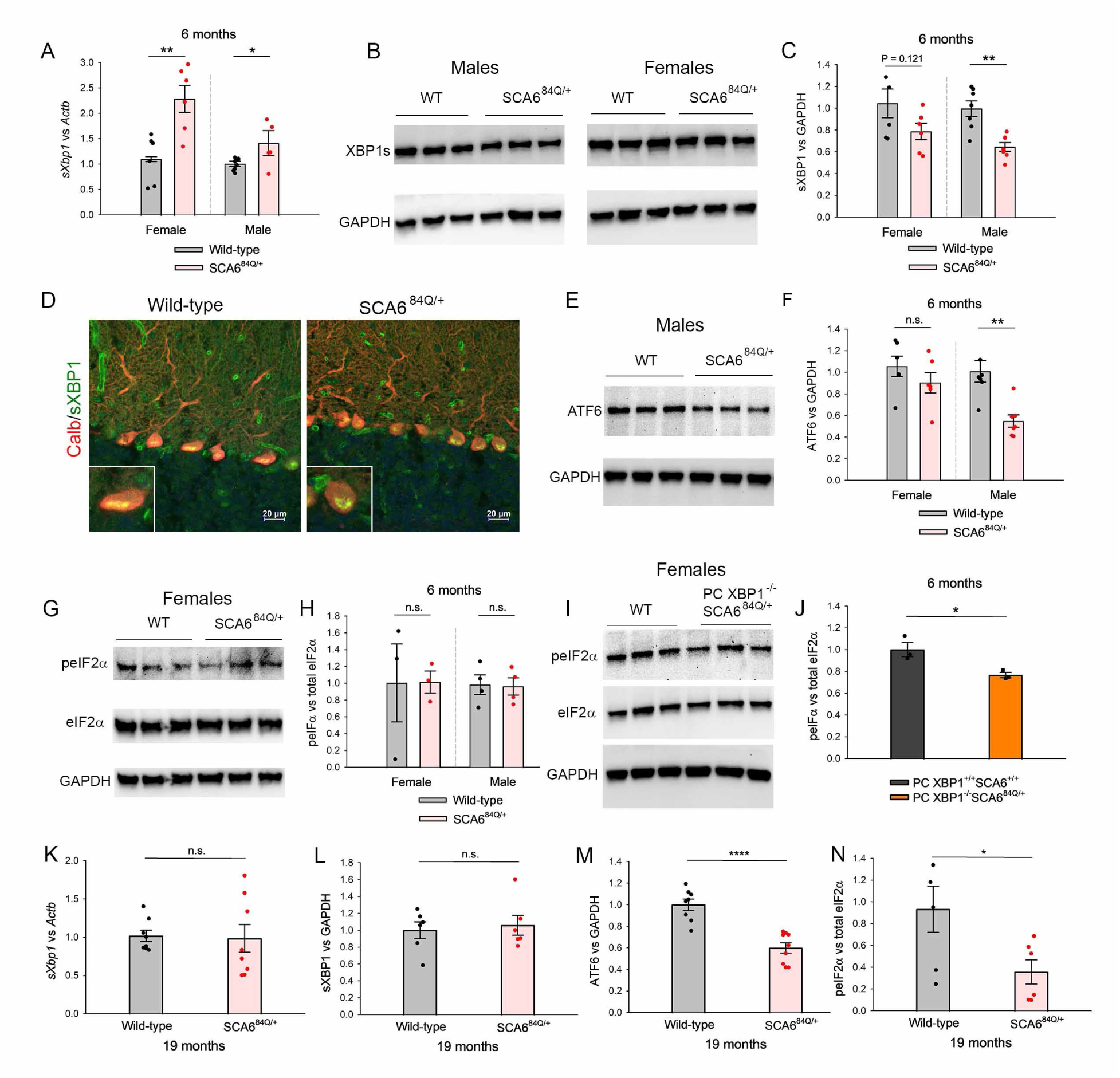
Alterations in all three canonical UPR pathways are present in SCA6. A) Quantitative RT-PCR of *sXbp1* in 6-month WT (N = 16) and SCA6^84Q/+^ mice (N = 15). The transcript levels of *sXbp1* are significantly increased in both 6-month male and female SCA6^84Q/+^ mice. Student’s t-test, *P < 0.05, **P < 0.01. B) Representative western blot showing reduced cerebellar sXBP1 levels in 6-month male and female SCA6^84Q/+^ mice and age- and sex-matched WT mice. C) Quantification of cerebellar sXBP1 normalized to cerebellar GAPDH levels in 6-month male (N =7) and female (N = 6) SCA6^84Q/+^ mice compared to WT male (N =7) and female (N =6) mice. Cerebellar sXBP1 levels are reduced in both male and female SCA6^84Q/+^ mice, although the reduction is only significant in male mice. Student’s t-test, **P < 0.01. D) Immunostaining showing sXBP1 (green) and Calbindin (red) in 6-month WT and SCA6^84Q/+^ mice. Note overall reduced staining of sXBP1 in Purkinje neuron nuclei in SCA6^84Q/+^ mice. Inset of WT Purkinje neuron (cell #3 from right) and SCA6^84Q/+^ Purkinje neuron (cell #1 from right) showing that in some SCA6^84Q/+^ Purkinje neurons, sXBP1 appears to form distinct puncta. E) Representative western blot showing reduced cerebellar ATF6 levels in 6-month male SCA6^84Q/+^ mice compared to 6-month WT male mice. F) Quantification of cerebellar ATF6 levels normalized to cerebellar GAPDH levels in 6-month male (N =7) and female (N = 6) SCA6^84Q/+^ mice compared to WT male (N =7) and female (N =6) mice. Cerebellar ATF6 levels are reduced only in 6-month male but not female SCA6^84Q/+^ mice. Student’s t-test, **P < 0.01, n.s.: Not significant. G) Representative western blot showing comparable cerebellar levels of peIF2α/eIF2α in 6-month female WT mice and female SCA6^84Q/+^ mice. H) Quantification of the ratio of cerebellar peIF2α to eIF2α levels, each separately normalized to cerebellar GAPDH levels, in 6-month male (N = 4) and female (N = 3) SCA6^84Q/+^ mice compared to WT male (N = 4) and female (N = 3) mice. Cerebellar peIF2α/eIF2α levels are unchanged in both 6-month male and female SCA6^84Q/+^ mice. Student’s t-test, n.s.: Not significant. I) Representative western blot showing reduced cerebellar peIF2α/eIF2α levels in 6-month female PC XBP1^-/-^SCA6^84Q/+^ mice compared to 6-month female PC XBP1^+/+^SCA6^+/+^ mice J) Quantification of cerebellar peIF2α/eIF2α levels in 6-month female PC XBP1^-/-^ SCA6^84Q/+^ mice (N = 3) compared to 6-month female PC XBP1^+/+^SCA6^+/+^ mice (N = 3). Cerebellar peIF2α/eIF2α levels are significantly reduced in female PC XBP1^-/-^SCA6^84Q/+^ mice. Student’s t-test, *P < 0.05. K) Quantitative RT-PCR of *sXbp1* in 19-month WT (N = 8) and SCA6^84Q/+^ mice (N = 8). The transcript levels of *sXbp1* are comparable between WT and SCA6^84Q/+^ mice. Student’s t-test, n.s.: Not significant. L) Quantification of cerebellar sXBP1 levels in 19-month WT (N = 6) and SCA6^84Q/+^ (N =6) mice. Cerebellar sXBP1 levels are similar between 19-month WT and SCA6^84Q/+^ mice. Student’s t-test, n.s.: Not significant. M) Quantification of cerebellar ATF6 levels in 19-month WT (N = 8) and SCA6^84Q/+^ (N =9) mice. Cerebellar ATF6 levels are significantly reduced in 19-month SCA6^84Q/+^ mice. Student’s t-test, ****P < 0.0001. N) Quantification of cerebellar peIF2α/eIF2α levels in 19-month WT (N = 6) and SCA6^84Q/+^(N =6) mice. Cerebellar peIF2α/eIF2α levels are significantly reduced in 19-month SCA6^84Q/+^ mice. Student’s t-test, *P < 0.05.

To examine the role for XBP1 in SCA6^84Q/+^ Purkinje neurons further, we generated SCA6^84Q/+^ mice where XBP1 is deleted in cerebellar Purkinje neurons (Figure S4B). We obtained WT mice with XBP1 in Purkinje neurons (PC XBP1^+/+^SCA6^+/+^), WT mice without XBP1 in Purkinje neurons (PC XBP1^-/-^SCA6^+/+^), SCA6^84Q/+^ mice with XBP1 in Purkinje neurons (PC XBP1^+/+^SCA6^84Q/+^), and SCA6^84Q/+^ mice without XBP1 in Purkinje neurons (PC XBP1^-/-^SCA6^84Q/+^). We first evaluated motor performance in all 4 genotypes using the rotarod at 1-month and 6-months of age. We expected PC XBP1^-/-^SCA6^84Q/+^ mice to show motor impairment similar to 19-month SCA6^84Q/+^ mice (Figure S4C). We did not, however, observe motor impairment in PC XBP1^-/-^SCA6^84Q/+^ mice either at 1-month or 6-months of age (Figure S4D). We then examined Purkinje neuron spiking regularity and firing frequency in all 4 genotypes. Consistent with the lack of a motor phenotype, Purkinje neuron spiking regularity and firing frequency were comparable across all genotypes (Figure S4E,F). These results suggest that XBP1 alone does not mediate the resilience to ER stress in SCA6^84Q/+^ Purkinje neurons and that redundancy in UPR pathways is present. Since there was a more prominent increase in *sXbp1* transcript levels in female SCA6^84Q/+^ mice, we examined protein levels of XBP1, ATF6, and peIF2α/eIF2α in 6-month female PC XBP1^-/-^ SCA6^84Q/+^ mice. Interestingly, cerebellar peIF2α/eIF2α levels were significantly downregulated in 6-month female PC XBP1^-/-^SCA6^84Q/+^ mice (Figure 6I,J). Additionally, cerebellar sXBP1 and ATF6 levels were reduced, although the reduction was not statistically significant (Figure S4G,H).

Our transcriptome data indicated a lack of upregulation of UPR-related gene transcripts in 19-month SCA6^84Q/+^ mice. We therefore investigated changes in XBP1, ATF6, and peIF2α/eIF2α in 19-month SCA6^84Q/+^ mice. *sXbp1* transcript levels were comparable between 19-month WT and SCA6^84Q/+^ mice (Figure 4K). We separately examined UPR pathways in 19-month male and female SCA6^84Q/+^ mice. Since no sex differences were observed, we pooled 19-month male and female SCA6^84Q/+^ cerebellar samples together for analysis. Cerebellar sXBP1 levels were similar between 19-month WT and SCA6^84Q/+^ mice (Figure 4L). Cerebellar levels of both ATF6 and peIF2α/eIF2α were reduced in 19-month SCA6^84Q/+^ mice (Figure 4M,N). These results suggest that UPR-related pathways are engaged at all ages in SCA6 mice, indicating that ER stress is present across the lifespan of SCA6 mice.

### ER stress induces irregular Purkinje neuron spiking in SCA6

We wished to examine the relationship between the observed changes in Purkinje neuron spiking and upregulation of ER stress pathways in the cerebella of SCA6^84Q/+^ mice. The UPR is triggered by ER stress, and a potent cause of ER stress is ER calcium depletion (36, 37). Depletion of ER calcium activates plasma membrane calcium-release activated calcium (CRAC) channels (38). We therefore reasoned that ER calcium dyshomeostasis may be responsible for irregular Purkinje neuron spiking through the activation of CRAC channels in 19-month SCA6^84Q/+^ mice. We performed noninvasive cell-attached recordings in 19-month SCA6^84Q/+^ mice, and examined changes in spike regularity following perfusion of 20 μM YM 58483 (also known as BTP2), a canonical CRAC channel inhibitor (38), in acute cerebellar slices. Spiking regularity and firing frequency were examined before YM 58483 perfusion (Figure 5A) and 5 min following YM 58483 perfusion (Figure 5B). Inhibiting the CRAC current in Purkinje neurons from 19-month SCA6^84Q/+^ mice significantly improved spiking irregularity (Figure 5C) and increased firing frequency (Figure 5D). To examine whether the effects were SCA6^84Q/+^ specific, we perfused YM 58483 on acute cerebellar slices from 19-month WT mice. YM 58483 did not alter the spiking regularity (Figure 5E) or firing frequency (Figure 5F) in WT mice, indicating that an increase in CRAC current is specific to Purkinje neurons in 19-month SCA6^84Q/+^ mice. The CRAC channel subunit enriched in Purkinje neurons is the Orai2/Stim1 subunit containing channel complex (39). Quantitative RT-PCR revealed no changes in transcript levels of *Orai2* or *Stim1* in 3-month (Figure S5A), 12-month (Figure S5C), or 19-month (Figure S5D) SCA6^84Q/+^ mice, although a small increase of both transcripts was noted in 6-month SCA6^84Q/+^ mice (Figure S5B). No significant changes in transcripts of *Orai1/2/3* or *Stim1/2* were observed at any age in SCA6^84Q/+^ mice in the RNAseq dataset (Table S2). YM 58483 inhibits TRPC3 channels in addition to CRAC channels, and TRPC3 channels have been reported to contribute to store-operated calcium entry in Purkinje neurons (40). We, therefore, examined whether Pyr10, a TRPC3 inhibitor (41) could improve spiking irregularity in 19-month SCA6^84Q/+^ mice. Pyr10 failed to improve spiking regularity (Figure 5G) or change firing frequency (Figure 5H) in 19-month SCA6^84Q/+^ Purkinje neurons, indicating that the observed increase in Purkinje neuron spiking irregularity is likely largely due to an increased CRAC current.

**Figure 5.**
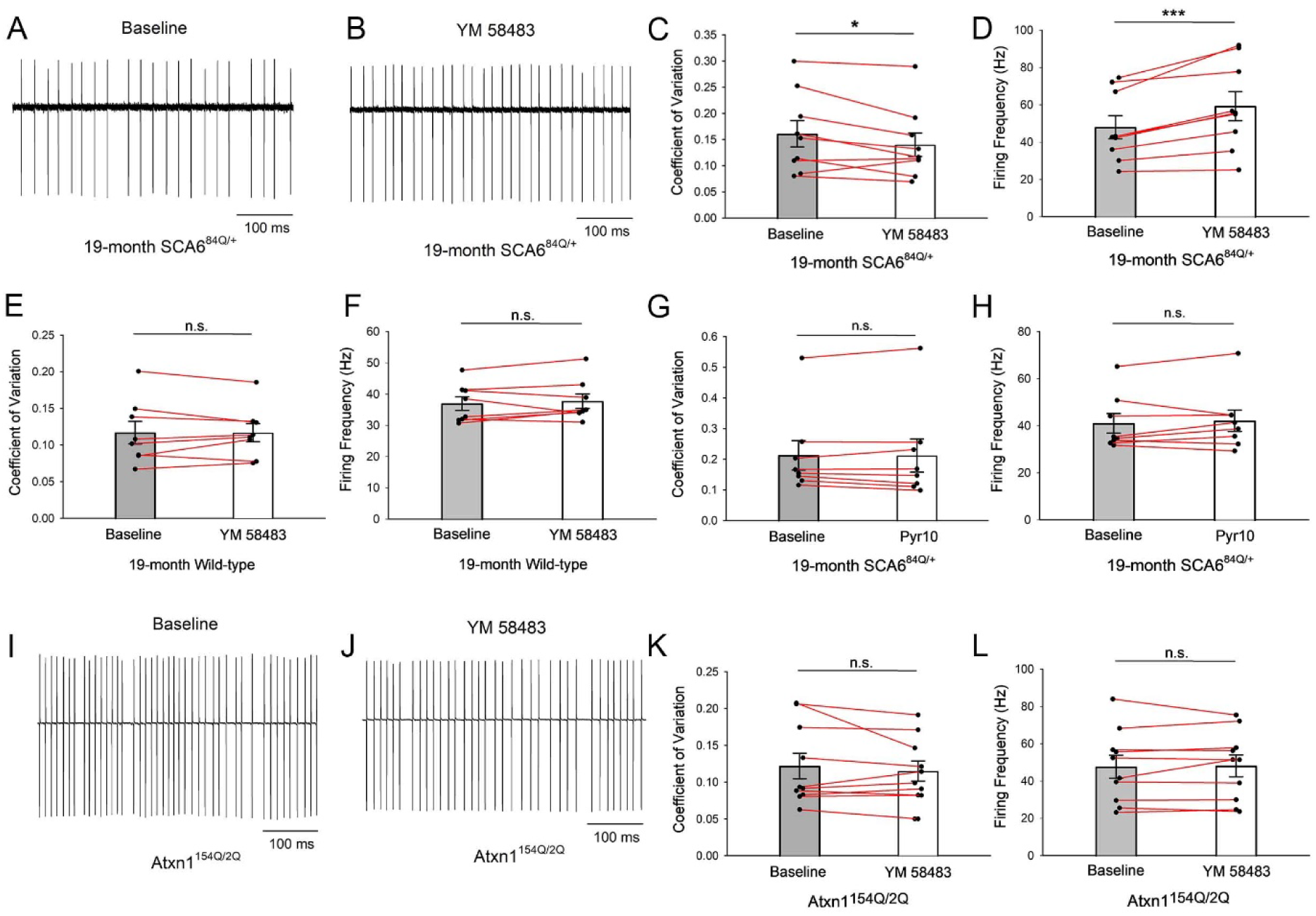
Irregular Purkinje neuron spiking in 19-month SCA6^84Q/+^ mice results from age-dependent activation of CRAC channels. A) Representative trace of baseline spontaneous Purkinje neuron irregular spiking in 19-month SCA6^84Q/+^ mice before 20 μM YM 58483 perfusion. B) Following perfusion of the CRAC channel inhibitor in the cell shown in 3A, spontaneous Purkinje neuron spiking irregularity improves in 19-month SCA6^84Q/+^ mice. C) YM 58483 significantly improves Purkinje neuron spiking irregularity in 19-month SCA6^84Q/+^ mice. n = 9 cells. Paired t-test, *P < 0.05. D) YM 58483 significantly increases Purkinje neuron firing frequency in 19-month SCA6^84Q/+^ mice. n = 9 cells. Paired t-test, ***P < 0.001. E) YM 58483 has no effect on Purkinje neuron spiking regularity in 19-month WT mice. n = 8 cells. Paired t-test, n.s.: Not significant. F) YM 58483 has no effect on Purkinje neuron firing frequency in 19-month WT mice. n = 8 cells. Paired t-test, n.s.: Not significant. G) Pyr10 has no effect on Purkinje neuron spiking regularity in 19-month SCA6^84Q/+^ mice. n = 8 cells. Paired t-test, n.s.: Not significant H) Pyr10 has no effect on Purkinje neuron firing frequency in 19-month SCA6^84Q/+^ mice. n = 8 cells. Paired t-test, n.s.: Not significant. I) Representative trace of baseline spontaneous irregular Purkinje neuron spiking in 14-week Atxn1^154Q/2Q^ mice before YM 58483 perfusion. J) Following perfusion of YM 58483 perfusion in the cell in 3I, there is no change in spiking irregularity in 14-week Atxn1^154Q/2Q^ mice. K) YM 58483 does not improve Purkinje neuron spiking irregularity in 14-week Atxn1^154Q/2Q^ mice. n = 10 cells. Paired t-test, n.s.: Not significant. L) YM 58483 has no effect on Purkinje neuron firing frequency in 14-week Atxn1^154Q/2Q^ mice. n = 10 cells. Paired t-test, n.s.: Not significant.

To investigate whether irregular Purkinje neuron spiking as a result of an increased CRAC current is unique to SCA6^84Q/+^ mice, we perfused YM 58483 on acute cerebellar slices from 14-week SCA1 mice (Atxn1^154Q/2Q^ mice), where Purkinje neuron spiking irregularity is similar to SCA6^84Q/+^ mice (28). Inhibiting CRAC had no effect on Purkinje neuron spiking regularity or firing frequency (Figure 5I-L), which is consistent with our prior studies showing that the irregular spiking in SCA1 is due to a reduced AHP and potassium channel transcripts (28). Taken together, our results indicate that an increase in CRAC current is the cause of the spiking irregularity in Purkinje neurons of symptomatic SCA6^84Q/+^ mice.

### HSP90 mitigates ER stress-induced membrane hyperexcitability

We wished to examine the mechanism for UPR-mediated compensation that prevents CRAC channel activation in SCA6^84Q/+^ Purkinje neurons. Of the many molecular chaperones whose transcripts are upregulated, we identified HSP90 as particularly important. HSP90 has been previously implicated in other late-onset neurodegenerative diseases characterized by protein aggregation (42), and upregulation of the ER-resident HSP90 isoform, GRP94 (encoded by *Hsp90b1*) is considered a hallmark of the UPR (43). We observed upregulation of transcripts of *Hsp90b1*, in addition to upregulated transcripts of *Hsp90aa1*, *Hsp90ab1* (cytosolic isoforms of HSP90), and the HSP90 co-chaperone *Ahsa1* (AHA1) in 6-month SCA6^84Q/+^ cerebella (Tables 1 and S2).

**Table 1.** Transcript levels of *Hsp90*-related genes are upregulated in 6-month SCA6^84Q/+^ mice.

| SCA6<sup>84Q/+</sup> x WT |  |  |  |  |  |  |
| --- | --- | --- | --- | --- | --- | --- |
| 6 Months |  |  |  |  |  |  |
| Gene | baseMean | log2FoldChange | lfcSE | stat | pvalue | padj |
| *Hsp90ab1* | 86001.45736 | 0.404713493 | 0.10649 | 3.80051 | 0.000144 | 0.003722 |
| *Hsp90b1* | 36971.0823 | 0.350081462 | 0.04772 | 7.33671 | 2.19E-13 | 1.10E-10 |
| *Hsp90aa1* | 74180.79037 | 0.331443086 | 0.04502 | 7.36153 | 1.82E-13 | 1.06E-10 |
| *Ahsa1* | 7838.91479 | 0.485988819 | 0.08152 | 5.96145 | 2.50E-09 | 4.55E-07 |

We investigated whether inhibiting HSP90 in 6-month presymptomatic SCA6^84Q/+^ mice could activate CRAC channels to produce the irregular Purkinje neuron spiking seen in symptomatic 19-month SCA6^84Q/+^ mice. We pre-incubated acute cerebellar slices in 1 μM 17-AAG, a pan-HSP90-specific reversible inhibitor (44) for an hour, to examine the role for HSP90 in maintaining regular spiking in 6-month SCA6^84Q/+^ Purkinje neurons. Purkinje neuron spiking was subsequently monitored non-invasively using cell-attached patch-clamp recordings in the presence of 1 μM 17-AAG. 17-AAG significantly increased Purkinje neuron spiking irregularity in 6-month SCA6^84Q/+^ mice (Figure 6A,B) but not in WT mice (Figures S6A,B). 17-AAG did not alter Purkinje neuron firing frequency in 6-month SCA6^84Q/+^ mice (Figure S6B). Importantly, perfusion of the CRAC current inhibitor, YM 58483, significantly improved the 17-AAG-induced irregular Purkinje neuron spiking in 6-month SCA6^84Q/+^ mice (Figure 6C) without altering firing frequency (Figure S6C). Additionally, YM 58483 did not produce a significant effect on Purkinje neuron spiking regularity (Figure 6D) or firing frequency (Figure S6D) in 17-AAG-pretreated 6-month WT cerebellar slices.

**Figure 6.**
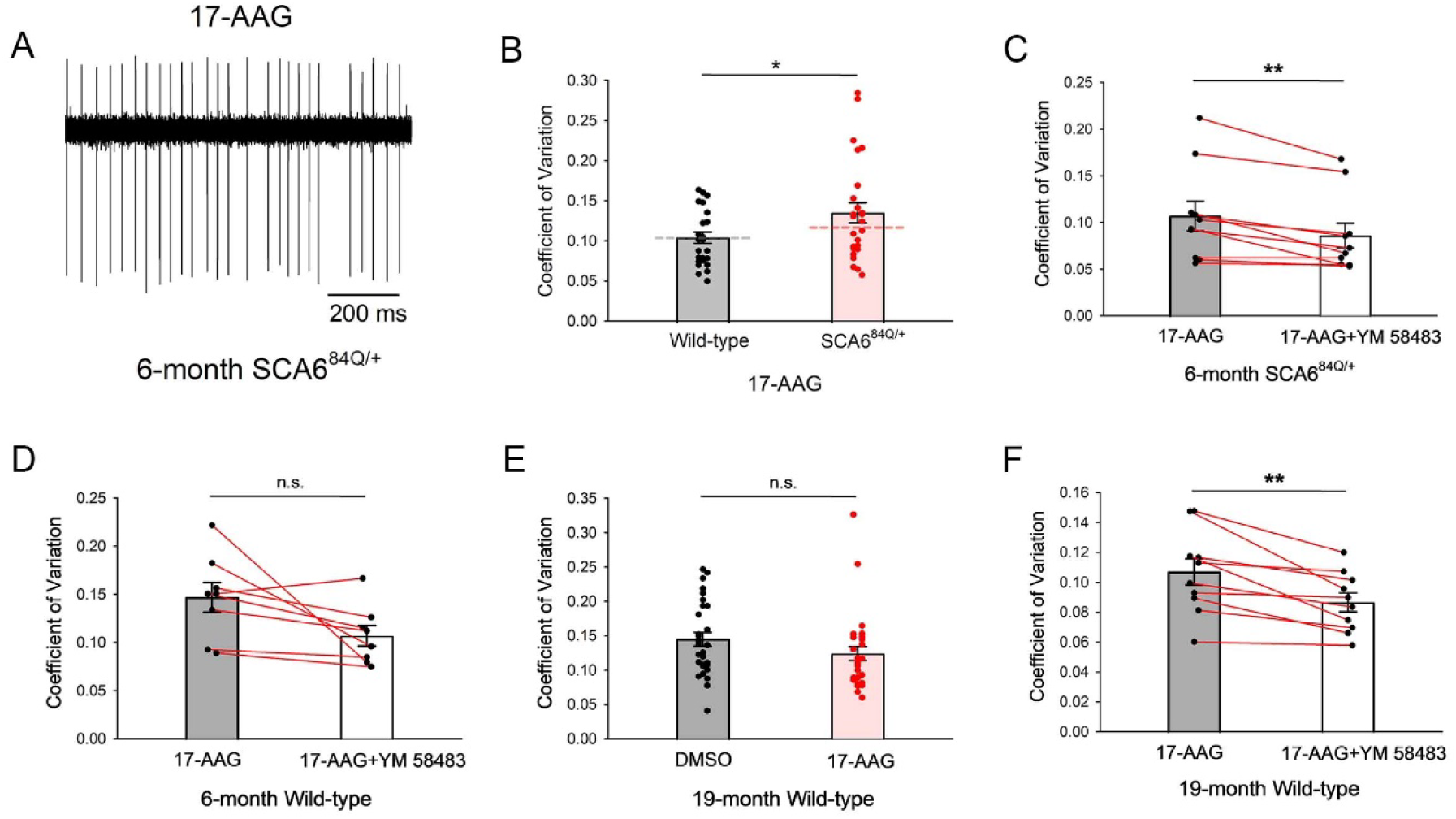
HSP90 mitigates ER stress-induced membrane hyperexcitability in pre- symptomatic SCA6^84Q/+^ mice. A) Representative trace of irregular Purkinje neuron spiking in 6-month SCA6^84Q/+^ cerebellar slices incubated for at least 1 hour with 1 μM 17-AAG, a pan-HSP90 reversible inhibitor. B) Pre-incubation with 1 μM 17-AAG has no effect on Purkinje neuron spiking regularity in 6-month WT mice. In 6-month SCA6^84Q/+^ cerebellar slices pre-incubated with 1 μM 17-AAG, Purkinje neuron spiking is irregular compared to WT mice. Dashed lines represent baseline Purkinje neuron spiking regularity from figure 1F. WT cells: n = 29, SCA6^84Q/+^ cells: n = 30. Student’s t-test, *P < 0.05. C) YM 58483 improves 17-AAG-induced Purkinje neuron spiking irregularity in 6- month SCA6^84Q/+^ mice. n = 10 cells. Paired t-test, **P < 0.01. D) YM 58483 has no effect on Purkinje neuron spiking in 17-AAG-pretreated cerebellar slices from 6-month WT mice. n = 8 cells. Paired t-test, n.s.: Not significant. E) 17-AAG has no effect on Purkinje neuron spiking regularity in 19-month WT mice. DMSO-treated cells: n = 29, 17-AAG-treated cells: n = 29. Student’s t-test, n.s.: Not significant. F) 17-AAG activates a CRAC current in 19-month WT Purkinje neurons as indicated by YM 58483 improving Purkinje neuron spiking regularity in 17-AAG-pretreated 19-month WT cerebellar slices. n = 10 cells. Paired t-test, **P < 0.01.

Next we sought to understand whether ER-stress related activation of CRAC channels occurs normally as a function of age. Inhibition of HSP90 with 17-AAG did not alter Purkinje neuron spiking regularity (Figure 6E) or firing frequency (Figure S6E) in 19-month WT mice. Unlike Purkinje neurons from 6-month WT mice, inhibition of the CRAC current significantly improved Purkinje neuron spiking regularity (Figure 6F) without changing firing frequency (Figure S6F) in 17-AAG treated 19-month WT cerebellar slices, indicating that 17-AAG was able to activate a CRAC current in19-month WT Purkinje neurons. These results suggest that there is an age-related compromise of the ability of HSP90 to mitigate ER-stress-related activation of CRAC channels, even though it does not reach the threshold, at least at 19 months of age, to cause abnormal Purkinje neuron spiking in WT mice. Taken together, our data indicate that activation of the UPR serves as a compensatory mechanism to mitigate ER-stress induced by polyQ Cav2.1 in SCA6 and that this resilience mechanism is compromised as a function of age, resulting in irregular Purkinje neuron spiking and disease onset.

## Discussion

A number of neurodegenerative disorders are associated with protein misfolding. It is, however, unclear whether aberrant proteostasis-induced ER stress is a driver of disease, or a target for therapeutic intervention. Our unbiased identification of a co-expressed gene network that is associated with resilience to ER-stress allows an interrogation of genes that are protective in late-onset neurodegenerative disease beyond SCA6. For example, examination of the network of genes in the lightcyan module identifies *Nrn1* expression as co-expressed with other ER-resilience genes regulated by XBP1, although *Nrn1* is not differentially expressed in SCA6^84Q/+^ mice. *Nrn1,* which encodes Neuritin, is a neurotrophic factor implicated in cognitive resilience in Alzheimer disease (45), but the mechanism through which it confers dendrite resilience to degeneration is unclear. Our data suggest that *Nrn1* may confer resilience to age related cognitive decline through mitigating ER-stress. These results also suggest that mitigation of ER stress may not be uniformly mediated by the same set of genes, and that resilience is conferred by genes that are unique to specific age-related neurodegenerative disorders.

The mechanism of disease in SCA6 is uncertain. When the CAG-repeat expansion in *CACNA1A* was initially identified as the cause of SCA6, and since SCA6 is allelic with episodic ataxia type 2, a childhood onset episodic disorder resulting from *CACNA1A* haploinsufficiency and without progressive neurodegeneration (4, 5), it was suggested that a loss of channel function could be responsible for disease. This was supported by *in vitro* studies in heterologous overexpression systems indicating that the biophysical properties of the channel were altered by the polyQ expansion in Cav2.1 (46–48). In Purkinje neurons of SCA6^84Q/+^ mice (7) and in human iPSC derived cortical neurons from individuals with SCA6 (49), however, the biophysical properties and current density of Cav2.1 are unchanged. Recent work has suggested that an alternative transcript of *CACNA1A*, α1ACT, residing in the C-terminus that contains the polyQ region, may contribute to disease in SCA6 (50). α1ACT acts as a transcription factor regulating genes involved in neurite outgrowth in early development (50). Viral-mediated expression of polyQ-α1ACT at postnatal day 1 results in motor impairment and Purkinje neuron degeneration by 4-postnatal weeks in mice (51). The early onset phenotype of polyQ-α1ACT transgenic mice is inconsistent with the late onset phenotype in human SCA6. SCA6^84Q/+^ mice don’t appear to have polyQ-α1ACT expression (7) and also more closely recapitulate the late onset of symptoms that is seen in human disease. A recent study suggests that in neural cells overexpressing either polyQ-α1ACT or polyQ-Cav2.1, mutant protein localizes to the ER, induces ER stress and mediates apoptosis in association with activation of the PERK-eIF2α pathway (52). This suggests that ER stress is primary mediator of disease in SCA6 rather than transcriptional dysregulation. The 84-repeat polyQ pathogenic expansion in SCA6^84Q/+^ mice, however, significantly exceeds the modest 21-30-polyQ pathogenic expansion in humans (2). The relevance of our identification of ER stress in SCA6^84Q/+^ should be interpreted cautiously in human SCA6 where the polyQ expansion is much more modest. A recent study revealed changes in mitochondrial structure and function following disease onset and during disease progression in SCA6^84Q/84Q^ mice (Leung et al., 2024). Homozygosity for the hyperexpanded polyQ repeat in Cav2.1 advances the age of onset to 7 months as compared to 19 months in heterozygous SCA6^84Q/+^ mice (7, 53). It is possible that the observed mitochondrial phenotypes are secondary to uncompensated ER stress due to the increased gene dosage of mutant *Cacna1a* in these mice. Future studies could examine the presence of ER stress in human iPSC derived neurons from individuals with SCA6.

Cav2.1 is expressed widely in the nervous system, and at particularly high levels in presynaptic terminals. It is therefore unclear why Purkinje neurons are uniquely vulnerable in SCA6. We have identified intact GABA_A_ transmission as necessary for ER stress-related Purkinje neuron calcium-dependent changes in membrane excitability in SCA6. Intracellular chloride homeostasis impacts ER stress and impacts neurodegeneration in cerebellar granule cells (54). Recent work has also indicated an important role for inhibitory transmission to Purkinje neurons in influencing Purkinje neuron degeneration in SCA1. Increased inhibitory transmission on Purkinje neurons was associated, paradoxically, with calcium-dependent Purkinje neuron dendritic hyperexcitability in SCA1 mice (55). The unique circuit properties in the cerebellum where molecular layer interneurons which are autonomously active, form synapses on Purkinje neuron somata, and provide tonic inhibitory input to Purkinje neurons (56), may make Purkinje neurons uniquely vulnerable to ER stress that is influenced by intracellular chloride homeostasis. Although we did not find changes in molecular layer interneuron-induced Purkinje neuron IPSC frequency in SCA6^84Q/+^ mice, in other polyQ ataxias such as SCA1 where there is an early increase in number and activity of molecular layer interneurons (55, 57), increased GABA_A_ inhibitory neurotransmission may be expected to compound the ER stress from expression of polyQ ATXN1 in Purkinje neurons.

Three canonical pathways are recognized to mitigate the deleterious consequences of ER stress. These are mediated by the XBP1, ATF6, and PERK-eIF2α pathways. UPR targets differ depending on cell type and context (20). Remarkably, the gene transcripts in the lightcyan module mediating the protective response in 6-month SCA6^84Q/+^ mice, including *Pdia4, Hsp90b1, Hspa5, Hyou1, Pdia6, Calr and Pdia3* overlap with gene transcripts that have been previously observed to be upregulated from enforced dimerization of sXBP1 and ATF6f (34). This enforced dimer of ATF6f/sXBP1 reduced the abnormal aggregation of mutant huntingtin and α-synuclein *in vitro* and demonstrated neuroprotection in preclinical models of Huntington’s and Parkinson’s disease. Our findings in SCA6 suggest that endogenous homeostatic mechanisms too can harness responses from more than one arm of the three canonical UPR pathways. We also identified redundancy the UPR pathways mitigating ER stress in SCA6. Deleting XBP1 specifically in cerebellar Purkinje neurons in SCA6^84Q/+^ mice resulted in downregulation of both ATF6 and peIF2α. Increased peIF2α has been linked to ER stress-induced apoptosis (26). Our results suggest that when XBP1-mediated upregulation of chaperone responses is abrogated, Purkinje neurons utilize alternative UPR pathways of downregulating peIF2α and ATF6. Additionally, our results suggest enhanced function of XBP1 in presymptomatic SCA6^84Q/+^ cerebella despite decreased protein levels. XBP1 has been shown to play competing roles in ER stress. XBP1 deficiency has been linked to enhanced macroautophagy and protects against Huntington’s disease (58). Enhanced XBP1 activity can promote protein folding, reduce protein aggregation and confer protection in other Huntington’s disease models (59). It is possible that XBP1 plays both roles simultaneously in SCA6^84Q/+^ mice to mediate resilience to ER stress. On the one hand, decreased levels of cerebellar XBP1 may promote degradation of polyQ-Cav2.1. On the other hand, enhanced function of XBP1 may simultaneously promote proper polyQ-Cav2.1 folding via upregulation of HSP90 and other chaperones. Further studies are needed to more fully understand the role of individual UPR pathways in SCA6.

Protein misfolding and changes in the UPR have been identified in a number of age-related neurodegenerative disorders, including in Parkinson’s, Alzheimer’s and Huntington’s disease (27). Calcium dyshomeostasis has also separately been implicated in these disorders (60–62). The identification of a mechanism of disease in SCA6 that connects abnormal proteostasis and calcium-dependent membrane excitability that also explains delayed onset of disease likely applies to a variety of other age-dependent neurodegenerative disorders. Exploring the relationship between aberrant proteostasis, ER stress and calcium dyshomeostasis further may be fruitful in late onset neurodegenerative disorders as enhancers of specific components of the UPR that confer protection from calcium dyshomeostasis could lead to novel treatment for SCA6 and other age-related neurodegenerative disorders.

## Materials and Methods

### Animals

All animal studies were reviewed and approved by the University of Texas Southwestern Medical Center. The SCA6^84Q/+^ mice were generated by the laboratory of Dr. Huda Y.Zoghbi (7) and deposited at the Jackson Lab. The SCA6^84Q/+^ mice used in the current study were obtained from the Jackson Laboratory through recovery of frozen embryos (RRID: IMSR_JAX:008683). The XBP1^flox/flox^ mice were a gift from Dr. Laurie Glimcher, who generated these mice (63). PCP2Cre^+/-^ mice (RRID:IMSR_JAX:010536), expressing cre recombinase solely in cerebellar Purkinje neurons (64) were obtained from Jackson Research Labs. PCP2Cre^+/-^ and SCA6^84Q/+^ mice in an XBP1^flox/flox^ background were first generated. PCP2Cre^+/-^XBP1^flox/flox^ and SCA6^84Q/+^ XBP1^flox/flox^ were crossed to subsequently generate PC XBP1^+/+^SCA6^+/+^ (XBP1^flox/flox^SCA6^+/+^ PCP2Cre^-/-^), PC XBP1^-/-^SCA6^+/+^ (XBP1^flox/flox^SCA6^+/+^ PCP2Cre^+/-^), PC XBP1^+/+^SCA6^84Q/+^ (XBP1^flox/flox^SCA6^84Q/+^ PCP2Cre^-/-^), and PC XBP1^-/-^SCA6^84Q/+^ (XBP1^flox/flox^SCA6^84Q/+^ PCP2Cre^+/-^) mice.

### Acute cerebellar slice preparation

Artificial CSF (aCSF) was prepared as follows (in mmol/L): 125 NaCl, 3.8 KCl, 26 NaHCO_3_, 1.25 NaH_2_PO_4_, 2 CaCl_2_, 1 MgCl_2_, 10 glucose. Mice were anesthetized with isoflurane inhalation before decapitation. Brains were rapidly removed for acute slices preparation. Slices were prepared at 300 µM on a VT1200 vibratome (Leica Biosystems, Buffalo Grove, IL) using the “hot cut” technique. aCSF were maintained between 33° and 36°C during tissue sectioning. Following sectioning, slices were incubated in pre-warmed, carbogen-bubbled (95% O_2_, 5% CO_2_) aCSF at 36°C for 45 min before recording. Slices were maintained in carbogen-bubbled aCSF at room temperature for the rest of the experiment.

### Patch clamp electrophysiology

Borosilicate glass patch pipettes of 3-5 MΩ were filled with an internal pipette solution containing (in mmol/L): 119 K-gluconate, 2 Na-gluconate, 6 NaCl, 2 MgCl_2_, 0.9 EGTA, 10 HEPES, 14 tris-phosphocreatine, 4 MgATP, 0.3 tris-GTP, at pH 7.3 and osmolarity 290 mOsm. Patch-clamp recordings were performed at 33°C. Pre-warmed, carbogen-bubbled aCSF was perfused at a rate of 150 mL/hour for all recordings. Recordings were acquired using an Axopatch 200B amplifier, Digidata 1440A interface, and pClamp-10 software (Molecular Devices, San Jose, CA). Current clamp data were acquired at 100 kHz in the fast current-clamp mode with bridge balance compensation, and filtered at 2 kHz. Cells were included only if the series resistance did not exceed 15 MΩ at any point during the recording, and if the series resistance did not change by more than 20% during the course of the recording. Voltage data have been corrected for a 10 mV liquid junction potential. A prior study in SCA6^84Q/+^ mice did not mention changes in spiking in relation to region of the cerebellum examined (8). Recordings were, therefore, performed without regard to anterior versus posterior cerebellum. The majority of the recordings were performed from the anterior vermis and all cells were pooled for analysis.

IPSC recordings were performed with an internal pipette solution containing (in mmol/L) 150 CsCl, 10 HEPES, 0.5 CaCl_2,_ 5 EGTA, at pH 7.2. Perforated patch clamp recordings used this same internal solution (33). Prior to perforated patch clamp IPSC recordings, gramicidin was dissolved in DMSO at a concentration of 2 mg/ml. The solution was then diluted in the electrode internal solution to obtain a final concentration of 5 µg/ml. Gramicidin containing electrode solution was made fresh every hour to ensure perforating potency. The solution was not filtered so as to not trap the antibiotic out of solution. The tip of the electrode was filled with regular internal solution to enhance seal formation. Input resistance was monitored through recording to ensure stable seal formation without significant leak. After obtaining baseline characterization in perforated patch mode, picrotoxin (50 µM) was perfused in the bath solution. The patch was then ruptured to enter conventional whole-cell recordings.

### Pharmacology

For some recordings, aCSF contained the following reagents (specified in the results section for individual experiments): YM 58483 (Tocris, cat#: 3939; 20 μM), 17-AAG (Cayman Chemicals, cat#: 11039; 1 μM), 6,7-dinitroquinoxaline-2,3-dione (DNQX) (Sigma Aldrich, cat#: D0540; 10 µM); picrotoxin (Sigma Aldrich, cat#: P1675; 50 µM). Purkinje neurons were recorded under baseline aCSF condition for at least 5 min before switching to reagent-contained aCSF. For 17- AAG in particular, slices were pre-incubated with 17-AAG containing aCSF for 1 hour at 36°C before recording. Slices were perfused with reagent-contained aCSF for at least 5 min and up to 10 min. All reported data were analyzed from the final 1 min of the baseline recording, and the final 1 min of the recording in the presence of the perfused compounds.

### RNA extraction and quantitative RT-PCR

Mice were deeply euthanized with isoflurane inhalation before decapitation. The cerebella were rapidly removed, flash-frozen in liquid nitrogen, and stored at −80°C for long-term use. RNA was isolated from the whole cerebellum using TRIzol (Life Technologies, cat#: 11596-026) and purified with RNeasy Mini Kit (Qiagen, cat#: 74104). Reverse transcription was performed with iScript cDNA Synthesis Kit (Bio-Rad, cat#: 1708891) using a C1000 Touch Thermal Cycler (Bio-Rad). Quantitative RT-PCR was performed with the iQ SYBR Green Supermix (Bio-Rad, cat#: 1708887) using a CFX96 Touch Real-Time PCR qPCR system (Bio-Rad). Gene expression was normalized to Actin (*Actb*) levels. Ct values for each gene were obtained in triplicate and averaged for statistical comparisons. Delta CT values were calculated as C ^target^ ^gene^ – C ^Actb^. Relative fold changes in gene expression were calculated using the 2^-ΔΔCt^ method (65). The primers (5’-3’) used for quantitative RT-PCR are listed below.

*Cacna1g* Forward: GTCGCTTTGGGTATCTTTGG

*Cacna1g* Reverse: TACTCCAGCATCCCAGCAAT

*Cacna1a* Forward: CCAGGAGCTCACCAAGGATG

*Cacna1a* Reverse: GCAGACAGGGGACTCACTTC

*Kcnma1* Forward: AGCCAACGATAAGCTGTGGT

*Kcnma1* Reverse: AATCTCAAGCCAAGCCAACT

*Itpr1* Forward: GGCAGAGATGATCAGGGAAA

*Itpr1* Reverse: AGCTCGTTCTGTTCCCCTTC

*Trpc3* Forward: GAGGTGAATGAAGGTGAACTGA

*Trpc3* Reverse: CGTCGCTTGGCTCTTATCTT

*Kcnn2* Forward: ACCCTAGTGGATCTGGCAAA

*Kcnn2* Reverse: GGGTG ACGATCCTTTTCTCA

*Orai2* Forward: ACAGTCAGGCCTGGTCC

*Orai2* Reverse: TGGTGGTTAGACGTGACGAG

*Stim1* Forward: CCTTGTCCATGCAGTCCCC

*Stim1* Reverse: GGGTCAAATCCCTCTGAGATCC

*sXbp1* Forward: GAGTCCGCAGCAGGTG

*sXbp1* Reverse: GTGTCAGAGTCCATGGGA

*Actb* Forward: CGGTTCCGATGCCCTGAGGCTCTT

*Actb* Reverse: CGTCACACTTCATGATGGAATTGA

### RNA sequencing and Weighted gene co-expression network analysis (WGCNA)

mRNA sequencing was performed on a NovaSeq instrument (Illumina, Inc.) with ∼155 million reads and 150 bp paired-end reads. 5 cerebellar samples from each genotype from each age were used. Samples were prepared individually and pooled together to reduce batch effects. The samples were then demultiplexed into fastq files and statistics were collected using FastQC (Babraham Bioinformatics). The fastq files were trimmed using Trimmomatic (66). 150 bp reads were aligned using STAR (67), and the percent of unique reads were calculated. Raw read counts per gene per sample were calculated using HTSeq (68). Outliers were removed based on principal component analysis and hierarchical clustering. Differential gene expression analysis was performed using DESeq2 (69). WGCNA was performed as described previously (70).

### Western blot analysis

Mice were deeply euthanized with isoflurane inhalation before decapitation. The cerebella were rapidly removed, flash-frozen in liquid nitrogen, and stored at −80°C for long-term use. Cerebellar tissue was homogenized using Teflon tissue homogenizer tube in Igepal lysis buffer (50 mM Tris-HCl, 150 mM NaCl, 5 mM EDTA, 1 mM EGTA, pH 8.0, 1% Igepal CA-640 (Sigma-Aldrich, cat#: 4906845001) with protease (Sigma-Aldrich, cat#: 4693132001) and phosphatase (Roche, cat# 04906837) inhibitor added. The whole cell lysates (WCL) were then centrifuged at 14000 rpm for 20 min at 4°C in temperature. The supernatant was then aliquoted and stored at −80°C. Lysates were subsequently thawed on ice, and protein concentration was determined using a Bicinchoninic acid (BCA) protein assay kit (Pierce BCA Protein Assay Kit, cat#: 23227). Samples were prepared for loading with 30 µg of protein denatured in 2X Laemmli Sample Buffer (Bio-Rad, cat#: 1610737) with 5% of 2-Mercaptoethanol (Sigma, cat#: M6250) at 95°C for 5 min. Protein samples were loaded on 10% precast gel (Bio-Rad, cat#: 4561034). The gel was run using Tris-Glycine-SDS buffer (Bio-Rad, cat#: 1610732) for 1.5 hours at a constant voltage of 100V and the resolved proteins were transferred on the polyvinylidene difluoride (PVDF) membrane, using Tris-Glycine buffer (Bio-Rad, cat#: 1610734) in 20% methanol for 1.5 to 2 hours at a constant current of voltage of 100V at 4°C. After transfer the membrane was blocked with 5% skimmed milk in TBS-T [Tris-buffered saline (TBS) and with tween 20 (T) (0.1% Tween 20)], for 1 hour at room temperature (RT). The membrane was then incubated with primary antibody overnight at 4°C according to the following dilutions in 5% BSA in TBS-T for immunoblot analysis: sXBP1 (Protein tech, cat#: 24868-1-AP) at 1:1000, ATF6 (Cell Signaling Technology, cat#: 65880) at 1:1000, phosphorylated eIF2α (Cell Signaling Technology, cat#: 3398) at 1:1000, eIF2α (Cell Signaling Technology, cat#: 5324) at 1:1000, GAPDH (Protein tech, cat#: 60004-I-Ig) at 1:5000 dilutions. After primary antibody incubation, the membrane was washed 3x10 min each with TBS-T and then incubated with secondary antibodies in 5% skimmed milk for 1 hour on rocker at RT. The secondary antibody was removed, and the membrane was washed 3x10 min each with TBS-T. The blots were developed using the Western blotting substrate (Thermo Scientific, cat#: 32209) under the Chemi-Doc imaging system (Bio-Rad) and the densitometry was performed using the Image Lab software from Bio-Rad.

### Tissue immunofluorescence and Microscopy

Mice were deeply euthanized with isoflurane inhalation before decapitation. Brains were placed in 1% paraformaldehyde for 1 hour at room temperature before transferring to a solution of 30% sucrose in phosphate-buffered saline (PBS) for 48 hours at 4°C. Brains were preserved in a 1:1 mixture of 30% sucrose in PBS:OCT compound (Fisher Scientific, cat#: 23-730-571) and stored at −80°C. Brains were sectioned to 20 µm thickness using a CM1850 cryostat (Leica). Tissue was permeabilized with 0.4% triton in PBS and blocked with 5% goat serum in 0.1% triton in PBS. Tissue was incubated with primary antibodies in PBS containing 0.1% triton and 2% normal goat serum at 4°C overnight. The following primary antibodies were used for immunostaining: rabbit anti-Cav2.1 (1:1000; Abcam, cat#: ab32642-200), mouse anti-Cav3.1 (1:150; NeuroMab, cat#: 75-206), sXBP1(1:400; Cell Signaling Technology, cat#: 40435), mouse anti-Calbindin (1:400; Sigma-Aldrich, cat#: C9848), rabbit anti-Calbindin (1:400; Sigma-Aldrich, cat#: C2724). Secondary antibody in PBS was applied to tissue for an hour at room temperature. Secondary antibodies used were as follows: Alexa Fluor 488 goat anti-mouse IgG (H+L) (1:200; Invitrogen, cat#: A11001), Alexa Fluor 488 goat anti-rabbit IgG (H+L) (1:200; Invitrogen, cat#: A11034), Alexa Fluor 594 goat anti-rabbit IgG (H+L) (1:200; Invitrogen, cat#: A11012), Alexa Fluor 594 goat anti-mouse IgG (H+L) (1:200; Invitrogen, cat#: A11032). Sections were imaged using an Axioskop 2 plus microscope (Zeiss). Cav2.1 and Cav3.1 intensity were quantified using ImageJ by measuring the mean pixel intensity within a box covering the molecular layer of cerebellar lobule 5. sXBP1 intensity was quantified using ImageJ by measuring the mean pixel intensity within a circle covering the nuclear region of Purkinje neurons. Representative images were acquired on a C2+ confocal microscope (Nikon) at 60x magnification.

### Statistical analysis

Data were compiled in Excel and analyzed using SigmaPlot. Electrophysiology data were analyzed using either unpaired or paired Student’s t-test as appropriate. Statistical significance was determined with an α-level of 0.05 for all studies. For RNAseq analysis, criteria used for filtering genes were an absolute log2 fold change ≥ 0.3 and an adjusted p-value ≤ 0.05.

### Data and Code Availability

The transcriptome dataset obtained from RNAseq analysis of cerebellar RNA isolated from 3, 6, 12, and 19-month WT and SCA6^84Q/+^ mice is available at the NCBI GEO website, accession GEO: GSE264100 (https://www.ncbi.nlm.nih.gov/geo/query/acc.cgi?acc=GSE264100). Code information can be found on GitHub (https://github.com/konopkalab/understanding_resilience_in_sca6).

## Acknowledgments

We are grateful to Dr. Laurie Glimcher (Dana Farber Cancer Institute) for Xbp1^flox/flox^ transgenic mice. This work was supported by the NIH R01 NS085054, and Raynor Cerebellum Project 23001199-SWMF to V.G.S..

## Author Contributions

HH, TLC, and VGS designed the research studies. HH, TLC, MF, MD, DMJ, and PK conducted experiments and acquired data. HH, TLC, MF, MD, DMJ, PK, AK, GK, and VGS analyzed data. HH and VGS wrote the manuscript.

