## Supplemental figures for "Resilience to Endoplasmic Reticulum Stress Mitigates Calcium-Dependent Membrane Hyperexcitability Underlying Late Disease Onset in SCA6"

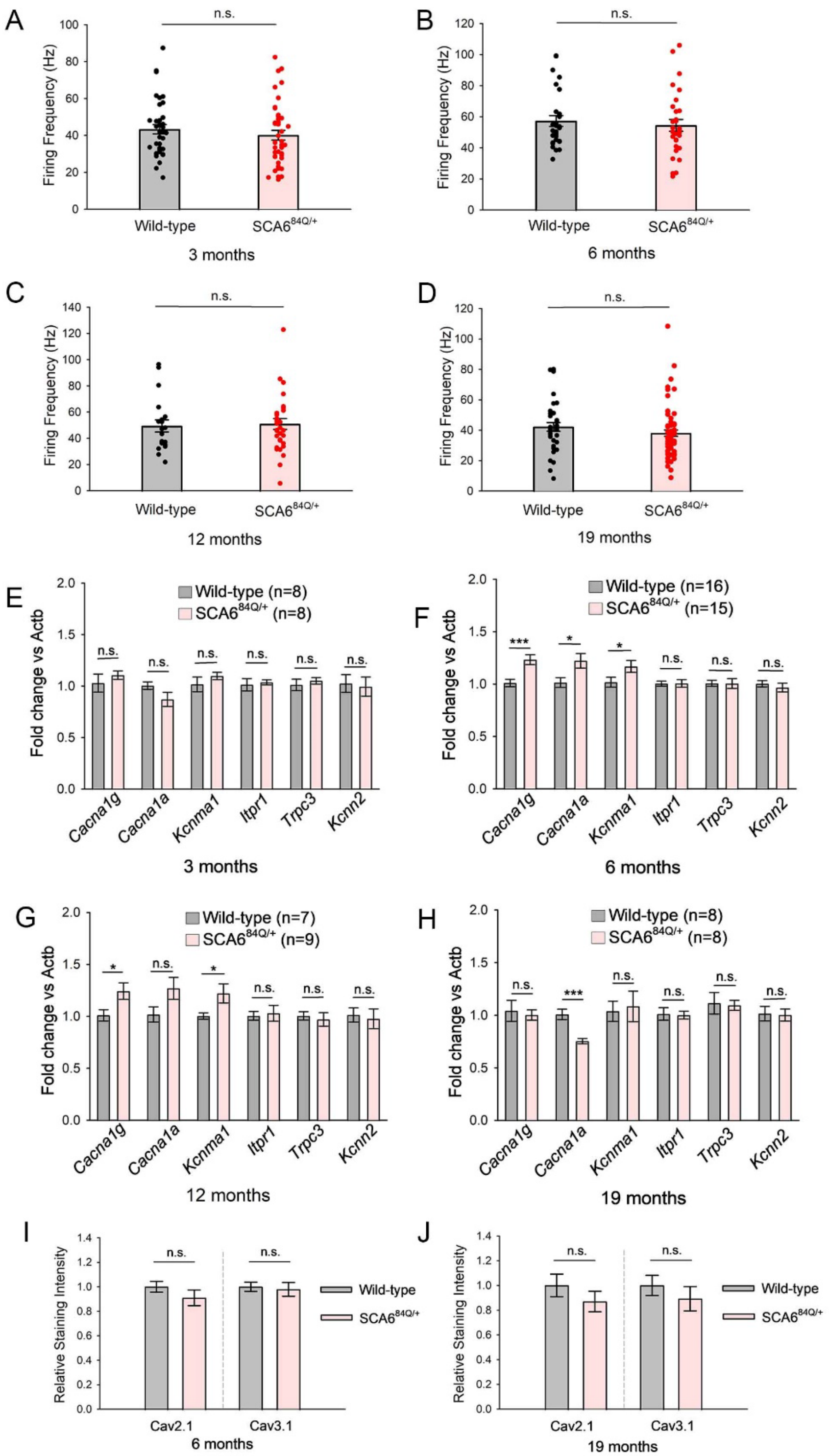

A
n.s.
Firing Frequency (Hz)
0
20
40
60
80
100
Wild-type
SCA6$^{84Q/+}$
3 months
B
n.s.
Firing Frequency (Hz)
0
20
40
60
80
100
120
Wild-type
SCA6$^{84Q/+}$
6 months
C
n.s.
Firing Frequency (Hz)
0
20
40
60
80
100
120
140
Wild-type
SCA6$^{84Q/+}$
12 months
D
n.s.
Firing Frequency (Hz)
0
20
40
60
80
100
120
Wild-type
SCA6$^{84Q/+}$
19 months
E
Wild-type (n=8)
SCA6$^{84Q/+}$ (n=8)
Fold change vs Actb
0.0
0.5
1.0
1.5
2.0
n.s.
n.s.
n.s.
n.s.
n.s.
n.s.
*Cacna1g*
*Cacna1a*
*Kcnma1*
*Itpr1*
*Trpc3*
*Kcnn2*
3 months
F
Wild-type (n=16)
SCA6$^{84Q/+}$ (n=15)
Fold change vs Actb
0.0
0.5
1.0
1.5
2.0
***
*
*
n.s.
n.s.
n.s.
*Cacna1g*
*Cacna1a*
*Kcnma1*
*Itpr1*
*Trpc3*
*Kcnn2*
6 months
G
Wild-type (n=7)
SCA6$^{84Q/+}$ (n=9)
Fold change vs Actb
0.0
0.5
1.0
1.5
2.0
*
n.s.
*
n.s.
n.s.
n.s.
*Cacna1g*
*Cacna1a*
*Kcnma1*
*Itpr1*
*Trpc3*
*Kcnn2*
12 months
H
Wild-type (n=8)
SCA6$^{84Q/+}$ (n=8)
Fold change vs Actb
0.0
0.5
1.0
1.5
2.0
n.s.
***
n.s.
n.s.
n.s.
n.s.
*Cacna1g*
*Cacna1a*
*Kcnma1*
*Itpr1*
*Trpc3*
*Kcnn2*
19 months
I
Relative Staining Intensity
0.0
0.2
0.4
0.6
0.8
1.0
1.2
1.4
n.s.
n.s.
Cav2.1
Cav3.1
6 months
Wild-type
SCA6$^{84Q/+}$
J
Relative Staining Intensity
0.0
0.2
0.4
0.6
0.8
1.0
1.2
1.4
n.s.
n.s.
Cav2.1
Cav3.1
19 months
Wild-type
SCA6$^{84Q/+}$

**Supplemental Figure 1. Purkinje neuron firing frequency and changes in ion channel transcripts/protein in SCA6$^{84Q/+}$ mice. [Related to Figure 1]**

A) Purkinje neuron firing frequency is unchanged between 3-month WT (N = 3) and SCA6$^{84Q/+}$ mice (N = 3). WT cells: n = 37, SCA6$^{84Q/+}$ cells: n = 42. Student's t-test, n.s.: Not significant.

B) Purkinje neuron firing frequency is unchanged between 6-month WT (N = 3) and SCA6$^{84Q/+}$ mice (N = 3). WT cells: n=28, SCA6$^{84Q/+}$ cells: n=30. Student's t-test, n.s.: Not significant.

C) Purkinje neuron firing frequency is unchanged between 12-month WT (N = 4) and SCA6$^{84Q/+}$ mice (N = 3). WT cells: n=20, SCA6$^{84Q/+}$ cells: n=31. Student's t-test, n.s.: Not significant.

D) Purkinje neuron firing frequency is unchanged between 19-month WT (N = 4) and SCA6$^{84Q/+}$ mice (N = 11). WT cells: n=35, SCA6$^{84Q/+}$ cells: n=64. Student's t-test, n.s.: Not significant.

E) Quantitative RT-PCR showing changes in the transcript levels of *Cacna1g*, *Cacna1a*, *Kcnma1*, *Itpr1*, *Trpc3*, and *Kcnn2* relative to *Actb* in 3-month WT (N = 8) and SCA6$^{84Q/+}$ (N = 8) mice. Student's t-test, n.s.: Not significant.

F) Quantitative RT-PCR showing changes in the transcript level of *Cacna1g*, *Cacna1a*, *Kcnma1*, *Itpr1*, *Trpc3*, and *Kcnn2* relative to *Actb* in 6-month WT (N = 16) and SCA6$^{84Q/+}$ (N = 15) mice. Student's t-test, *P < 0.05, ***P < 0.001, n.s.: Not significant.

G) Quantitative RT-PCR showing changes in the transcript level of *Cacna1g*, *Cacna1a*, *Kcnma1*, *Itpr1*, *Trpc3*, and *Kcnn2* relative to *Actb* in 12-month WT (N = 7) and SCA6$^{84Q/+}$ (N = 9) mice. Student's t-test, *P < 0.05, n.s.: Not significant.

H) Quantitative RT-PCR showing changes in the transcript level of *Cacna1g*, *Cacna1a*, *Kcnma1*, *Itpr1*, *Trpc3*, and *Kcnn2* relative to *Actb* in 19-month WT (N = 8) and SCA6$^{84Q/+}$ (N = 8) mice. Student's t-test, ***P < 0.001, n.s.: Not significant.

I) Quantification of immunostaining showing that protein levels of Cav2.1 and Cav3.1 are unchanged between cerebella of 6-month WT (N = 4) and $SCA6^{84Q/+}$ (N = 4) mice. Student's t-test, n.s.: Not significant.

J) Quantification of immunostaining showing that protein levels of Cav2.1 and Cav3.1 are unchanged between cerebella of 19-month WT (N = 5) and $SCA6^{84Q/+}$ (N = 5) mice. Student's t-test, n.s.: Not significant.

### 6 months

| | Wide-type | SCA6$^{84Q/+}$ | P-value |
|---|---|---|---|
| **AHP Minimum (mV)** | -66.0190 ± 0.6839 | -66.0626 ± 0.5831 | 0.9622 |
| **Spike Peak (mV)** | 12.3835 ± 0.3196 | 13.3399 ± 0.5410 | 0.1279 |
| **Depolarizaton dV/dt (mV/ms)** | 585.4311 ± 21.4718 | 573.9865 ± 18.0998 | 0.6904 |
| **Repolarization dV/dt (mV/ms)** | -438.8379 ± 15.8392 | -445.0329 ± 15.58 | 0.7831 |
| **Spike Half-width (mV)** | 0.0791 ± 0.0028 | 0.0821 ± 0.0037 | 0.5115 |
| **Input Resistance (MΩ)** | 65.5695 ± 3.7851 | 65.6998 ± 4.7366 | 0.9828 |

### 19 months

| | Wide-type | SCA6$^{84Q/+}$ | P-value |
|---|---|---|---|
| **AHP Minimum (mV)** | -64.9970 ± 0.3509 | -64.5236 ± 0.5243 | 0.4389 |
| **Spike Peak (mV)** | 10.8325 ± 0.4774 | 13.9117 ± 0.3789 | 1.45E-05 |
| **Depolarizaton dV/dt (mV/ms)** | 383.6699 ± 11.2179 | 411.9101 ± 18.4150 | 0.1732 |
| **Repolarization dV/dt (mV/ms)** | -278.1233 ± 7.8184 | -302.3107 ± 13.5810 | 0.1069 |
| **Spike Half-width (mV)** | 0.0974 ± 0.0027 | 0.1138 ± 0.0046 | 0.0020 |
| **Input Resistance (MΩ)** | 79.2219 ± 3.4212 | 74.4427 ± 3.9312 | 0.3647 |

**Supplementary Table 1. Electrophysiological parameters of Purkinje neurons from 6-month and 19-month wild-type and SCA6$^{84Q/+}$ mice**.

Data are presented as mean ± standard error of the mean. Two-tailed Student’s t-test.

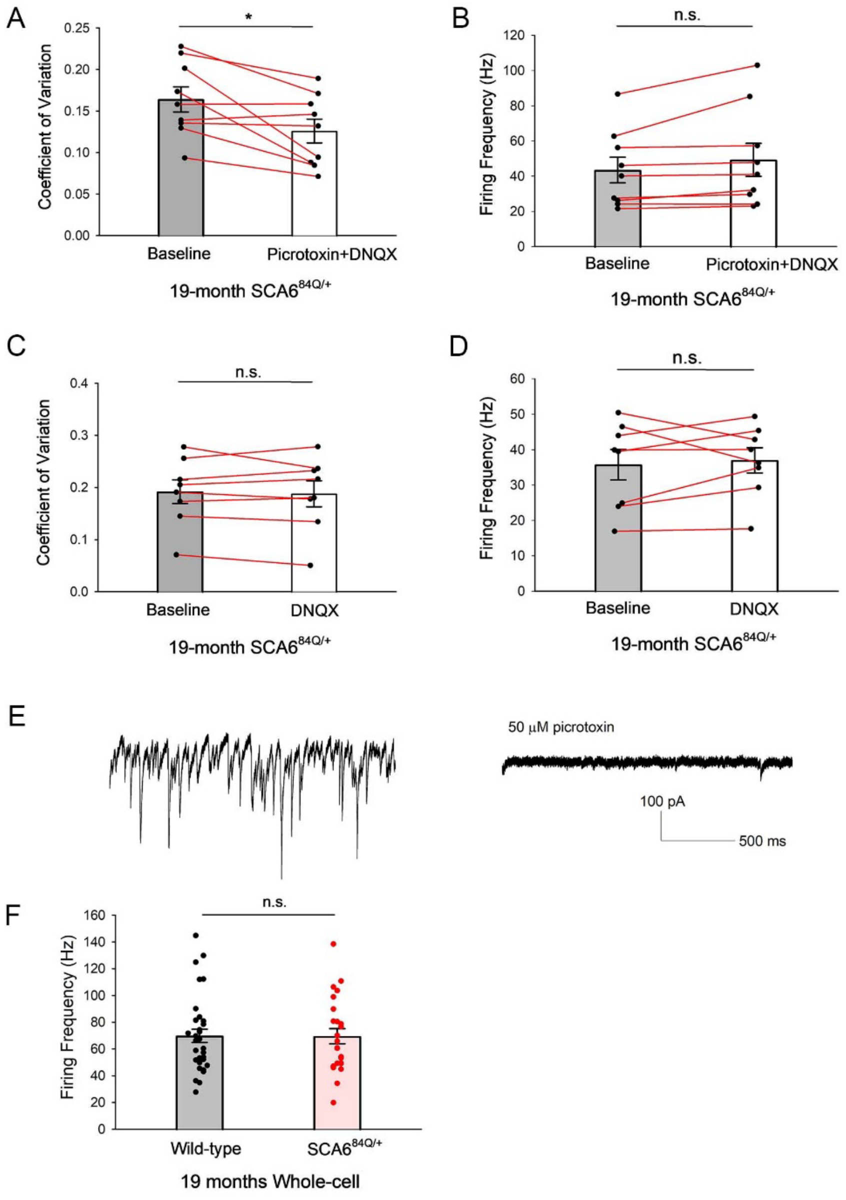

A
*
Coefficient of Variation
0.25
0.20
0.15
0.10
0.05
0.00
Baseline
Picrotoxin+DNQX
19-month SCA6^84Q/+
B
n.s.
Firing Frequency (Hz)
120
100
80
60
40
20
0
Baseline
Picrotoxin+DNQX
19-month SCA6^84Q/+
C
n.s.
Coefficient of Variation
0.4
0.3
0.2
0.1
0.0
Baseline
DNQX
19-month SCA6^84Q/+
D
n.s.
Firing Frequency (Hz)
60
50
40
30
20
10
0
Baseline
DNQX
19-month SCA6^84Q/+
E
50 μM picrotoxin
100 pA
500 ms
F
n.s.
Firing Frequency (Hz)
160
140
120
100
80
60
40
20
0
Wild-type
SCA6^84Q/+
19 months Whole-cell

**Supplemental Figure 2. Intact inhibitory transmission is necessary to activate Purkinje neuron CRAC channels in SCA6$^{84Q/+}$ mice. [Related to Figure 2]**

A) A combination of picrotoxin and DNQX improves Purkinje neuron spiking irregularity in 19-month SCA6$^{84Q/+}$ mice. n = 9 cells. Paired t-test, *P < 0.05.

B) A combination of picrotoxin and DNQX has no effect on Purkinje neuron firing frequency in 19-month SCA6$^{84Q/+}$ mice. n = 9 cells. Paired t-test, n.s.: Not significant.

C) DNQX alone has no effect on Purkinje neuron spiking regularity in 19-month SCA6$^{84Q/+}$ mice. n = 8 cells. Paired t-test, n.s.: Not significant.

D) DNQX alone has no effect on Purkinje neuron firing frequency in 19-month SCA6$^{84Q/+}$ mice. n = 8 cells. Paired t-test, n.s.: Not significant.

E) Representative traces of IPSCs before (left) and after (right) picrotoxin perfusion.

F) Purkinje neuron firing frequency in the whole-cell patch clamp configuration is comparable between 19-month WT and SCA6$^{84Q/+}$ mice. WT cells: n = 33, SCA6$^{84Q/+}$ cells: n = 24. Student’s t-test, n.s.: Not significant.

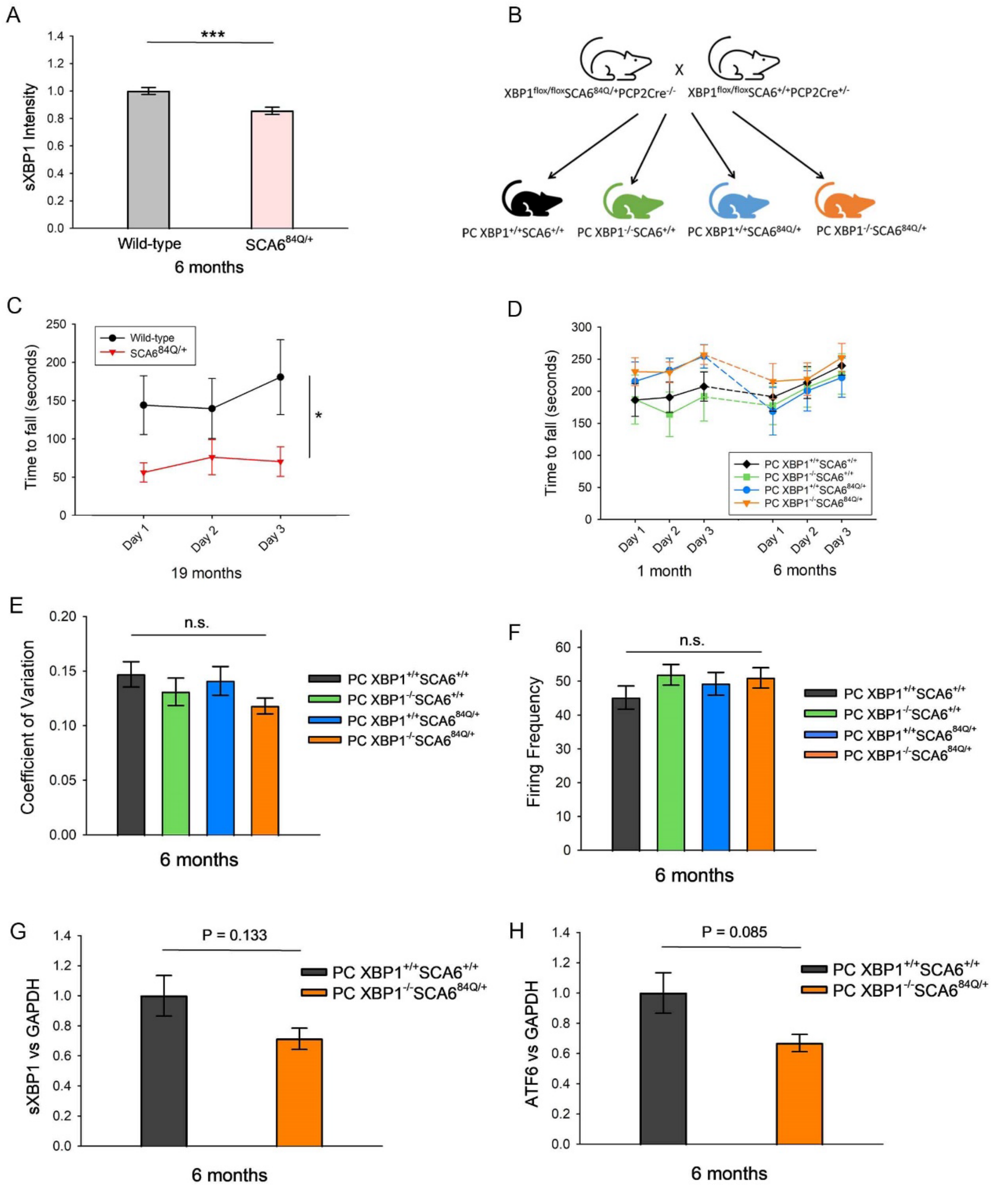
A
sXBP1 Intensity
1.4
1.2
1.0
0.8
0.6
0.4
0.2
0.0
***
Wild-type
SCA6^84Q/+
6 months
B
XBP1^flox/flox SCA6^84Q/+ PCP2Cre^-/-
X
XBP1^flox/flox SCA6^+/+ PCP2Cre^+/-
PC XBP1^+/+ SCA6^+/+
PC XBP1^-/- SCA6^+/+
PC XBP1^+/+ SCA6^84Q/+
PC XBP1^-/- SCA6^84Q/+
C
Time to fall (seconds)
250
200
150
100
50
0
Wild-type
SCA6^84Q/+
*
Day 1
Day 2
Day 3
19 months
D
Time to fall (seconds)
300
250
200
150
100
50
0
PC XBP1^+/+ SCA6^+/+
PC XBP1^-/- SCA6^+/+
PC XBP1^+/+ SCA6^84Q/+
PC XBP1^-/- SCA6^84Q/+
Day 1
Day 2
Day 3
Day 1
Day 2
Day 3
1 month
6 months
E
Coefficient of Variation
0.20
0.15
0.10
0.05
0.00
n.s.
PC XBP1^+/+ SCA6^+/+
PC XBP1^-/- SCA6^+/+
PC XBP1^+/+ SCA6^84Q/+
PC XBP1^-/- SCA6^84Q/+
6 months
F
Firing Frequency
60
50
40
30
20
10
0
n.s.
PC XBP1^+/+ SCA6^+/+
PC XBP1^-/- SCA6^+/+
PC XBP1^+/+ SCA6^84Q/+
PC XBP1^-/- SCA6^84Q/+
6 months
G
sXBP1 vs GAPDH
1.4
1.2
1.0
0.8
0.6
0.4
0.2
0.0
P = 0.133
PC XBP1^+/+ SCA6^+/+
PC XBP1^-/- SCA6^84Q/+
6 months
H
ATF6 vs GAPDH
1.4
1.2
1.0
0.8
0.6
0.4
0.2
0.0
P = 0.085
PC XBP1^+/+ SCA6^+/+
PC XBP1^-/- SCA6^84Q/+
6 months

**Supplemental Figure 4. Knockout of XBP1 in Purkinje neurons has no effect on the motor phenotype, Purkinje neuron spiking regularity and firing frequency but alters other UPR pathways in SCA6$^{84Q/+}$ mice. [Related to Figure 4]**

A) Quantification of sXBP1 intensity in nuclei of 6-month WT (n = 39 cells) and SCA6$^{84Q/+}$ (n = 65 cells) Purkinje neurons. The nuclear sXBP1 intensity is significantly reduced in SCA6$^{84Q/+}$ Purkinje neurons. Student's t-test, ***P < 0.001.

B) Mouse breeding scheme showing the generation of PC XBP1$^{+/+}$SCA6$^{+/+}$ (XBP1$^{flox/flox}$SCA6$^{+/+}$ PCP2Cre$^{-/-}$), PC XBP1$^{-/-}$SCA6$^{+/+}$ (XBP1$^{flox/flox}$SCA6$^{+/+}$ PCP2Cre$^{+/-}$), PC XBP1$^{+/+}$SCA6$^{84Q/+}$ (XBP1$^{flox/flox}$SCA6$^{84Q\ /+}$ PCP2Cre$^{-/-}$), and PC XBP1$^{-/-}$SCA6$^{84Q/+}$ (XBP1$^{flox/flox}$SCA6$^{84Q\ /+}$ PCP2Cre$^{+/-}$) mice.

C) 19-month SCA6$^{84Q/+}$ mice (N = 11) perform significantly worse on the rotarod compared to age-matched WT mice (N = 7). Student's t-test, *P < 0.05.

D) Motor performance on the rotarod is comparable among PC XBP1$^{+/+}$SCA6$^{+/+}$ mice (N = 15), PC XBP1$^{-/-}$SCA6$^{+/+}$ mice (N = 9), PC XBP1$^{+/+}$SCA6$^{84Q/+}$ mice (N = 7), and PC XBP1$^{-/-}$SCA6$^{84Q/+}$ mice (N = 12) at both 1-month and 6-months of age. Two-way repeated measures ANOVA.

E) Purkinje neuron spiking regularity, indicated by the coefficient of variation of the interspike interval, is comparable among PC XBP1$^{+/+}$SCA6$^{+/+}$ mice (n = 33 cells), PC XBP1$^{-/-}$SCA6$^{+/+}$ mice (n = 30 cells), PC XBP1$^{+/+}$SCA6$^{84Q/+}$ mice (n = 34 cells), and PC XBP1$^{-/-}$SCA6$^{84Q/+}$ mice (n = 34 cells) at 6 months of age. One way ANOVA, n.s.: Not significant.

F) Purkinje neuron firing frequency is unchanged among PC XBP1$^{+/+}$SCA6$^{+/+}$ mice (n = 33 cells), PC XBP1$^{-/-}$SCA6$^{+/+}$ mice (n = 30 cells), PC XBP1$^{+/+}$SCA6$^{84Q/+}$ mice (n = 34 cells), and PC XBP1$^{-/-}$SCA6$^{84Q/+}$ mice (n = 34 cells) at 6 months of age. One way ANOVA, n.s.: Not significant.

G) Quantification of cerebellar sXBP1 levels in female PC XBP1$^{-/-}$SCA6$^{84Q/+}$ mice (N = 3) compared to female PC XBP1$^{+/+}$SCA6$^{+/+}$ mice (N = 3). Cerebellar sXBP1 levels are reduced in female PC XBP1$^{-/-}$SCA6$^{84Q/+}$ mice but this reduction is not statistically significant. Student’s t-test.

H) Quantification of cerebellar ATF6 levels in female PC XBP1$^{-/-}$SCA6$^{84Q/+}$ mice (N = 3) compared to female PC XBP1$^{+/+}$SCA6$^{+/+}$ mice (N = 3). Cerebellar ATF6 levels are reduced in female PC XBP1$^{-/-}$SCA6$^{84Q/+}$ mice but this reduction is not statistically significant. Student’s t-test.

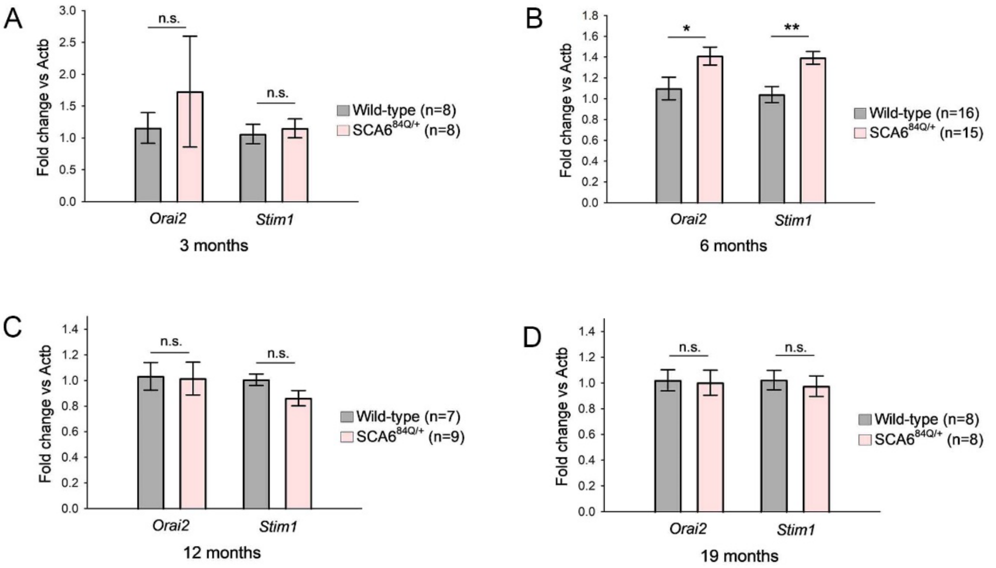


**Supplemental Figure 5. The increase in CRAC current in 19-month SCA6$^{84Q/+}$ mice is not caused by increased expression of Orai2/Stim1 subunits in cerebella. [Related to Figure 5]**

A) Quantitative RT-PCR shows no changes in *Orai2* or *Stim1* transcript levels relative to *Actb* in cerebella of 3-month SCA6$^{84Q/+}$ mice (N = 8) compared to WT mice (N = 8). Student's t-test, n.s.: Not significant.

B) Quantitative RT-PCR shows a small increase of both *Orai2* and *Stim1* transcripts relative to *Actb* in the cerebella of 6-month SCA6$^{84Q/+}$ mice (N = 15) compared to WT mice (N = 16). Student's t-test, *P < 0.05, **P < 0.01.

C) Quantitative RT-PCR shows no changes in *Orai2* or *Stim1* transcript levels relative to *Actb* in cerebella of 12-month SCA6$^{84Q/+}$ mice (N = 9) compared to WT mice (N = 7). Student's t-test, n.s.: Not significant.

D) Quantitative RT-PCR shows no changes in *Orai2* or *Stim1* transcript levels relative to *Actb* in cerebella of 19-month SCA6$^{84Q/+}$ mice (N = 8) compared to WT mice (N = 8). Student's t-test, n.s.: Not significant.

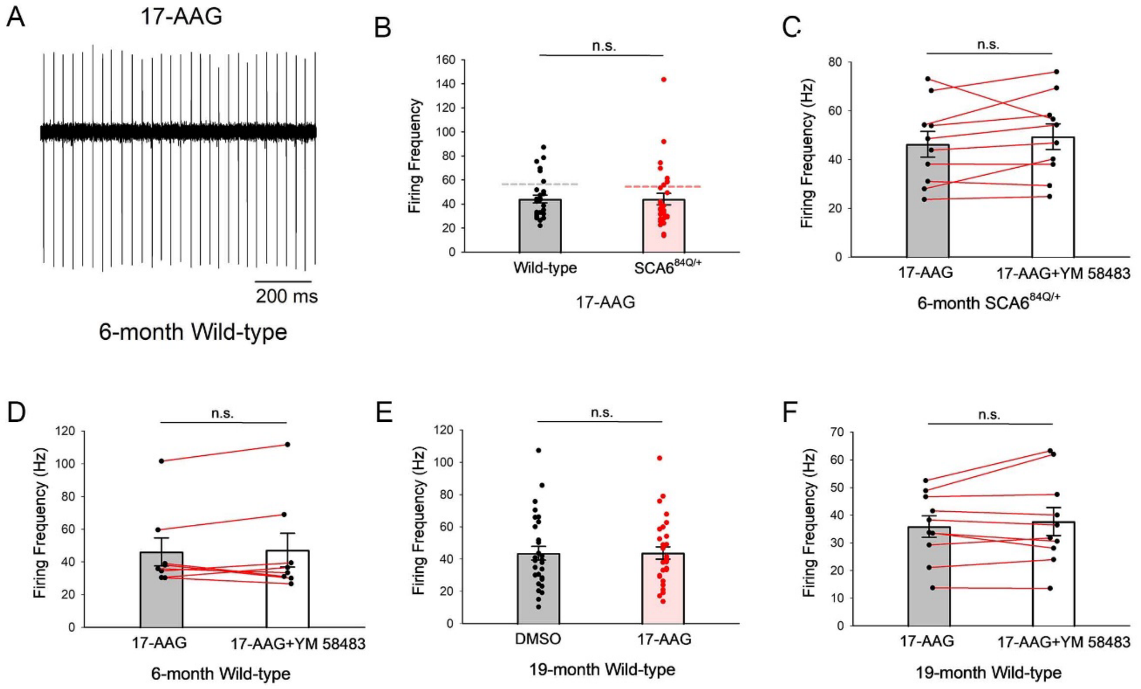


**Supplemental Figure 6. 17-AAG has no effect on Purkinje neuron firing frequency in 6-month SCA6$^{84Q/+}$ mice or 19-month wild-type mice. [Related to Figure 6]**

A) Representative trace of spontaneous Purkinje neuron spiking in 17-AAG pre-incubated cerebellar slices from 6-month WT mice.

B) 17-AAG does not change Purkinje neuron firing frequency in 6-month WT or SCA6$^{84Q/+}$ mice. Dashed lines represent the mean baseline firing frequency from Figure S1B. WT cells: n = 29, SCA6$^{84Q/+}$ cells: n = 30. Student's t-test, n.s.: Not significant.

C) YM 58483 does not alter firing frequency in 17-AAG-pretreated Purkinje neurons in 6-month SCA6$^{84Q/+}$ mice. n = 10 cells. Paired t-test, n.s.: Not significant.

D) YM 58483 does not alter firing frequency in 17-AAG-pretreated Purkinje neurons in 6-month WT mice. n = 8 cells. Paired t-test, n.s.: Not significant.

E) 17-AAG has no effect on Purkinje neuron firing frequency in 19-month WT mice. DMSO-treated cells: n = 29, 17-AAG-treated cells: n = 29. Student's t-test, n.s.: Not significant.

F) YM 58483 has no effect on Purkinje neuron firing frequency in 17-AAG-pretreated 19-month WT cerebellar slices. n = 10 cells. Paired t-test, n.s.: Not significant.
